# *BrainPy*: a flexible, integrative, efficient, and extensible framework towards general-purpose brain dynamics programming

**DOI:** 10.1101/2022.10.28.514024

**Authors:** Chaoming Wang, Xiaoyu Chen, Tianqiu Zhang, Si Wu

**Affiliations:** School of Psychology and Cognitive Sciences, IDG/McGovern Institute for Brain Research, Peking-Tsinghua Center for Life Sciences, Center of Quantitative Biology, Academy for Advanced Interdisciplinary Studies, Peking University, Beijing, 100871, China; Chinese Institute for Brain Research, Beijing, 100871, China

## Abstract

The neural mechanisms underlying brain functions are extremely complicated. Brain dynamics modeling is an indispensable tool for elucidating these mechanisms by modeling the dynamics of the neural circuits that execute brain functions. To ease and facilitate brain dynamics modeling, a general-purpose programming framework is needed to enable users to freely define neural models across multiple scales; efficiently simulate, train, and analyze model dynamics; and conveniently extend new modeling approaches. By utilizing the advanced just-in-time (JIT) compilation, we developed BrainPy. BrainPy provides a rich infrastructure tailored for brain dynamics programming, which supports an integrated platform for brain dynamics model building, simulation, training, and analysis. Models in BrainPy can be JIT compiled into binary instructions for multiple devices (including CPU, GPU, and TPU) to achieve a high running performance comparable to native C or CUDA. Moreover, BrainPy features an extensible architecture allowing easy expansion of new infrastructure, utilities, and machine learning approaches.

## 1 Introduction

Brain dynamics modeling, which uses computational models to simulate and elucidate brain functions, is receiving increasing attention from researchers across different disciplines. Recently, gigantic projects in brain science have been initiated worldwide, including the BRAIN Initiative [1], Human Brain Project [2], and China Brain Project [3], which are continuously producing new data about the structures and activity patterns of neural systems. Computational modeling is a fundamental and indispensable tool for interpreting this vast amount of data. However, to date, we still lack a general-purpose programming framework for brain dynamics modeling. By general-purpose, we mean that such a programming framework can implement most brain dynamics models, integrate diverse modeling demands (e.g., simulation, training, and analysis), accommodate new modeling approaches constantly emerging in the field while maintaining high running performance. An example of general-purpose programming is TensorFlow [4] or PyTorch [5] in the field of Deep Learning, which provides convenient interfaces for researchers to define various AI models flexibly and efficiently. These frameworks have become essential infrastructure in AI research, and play an indispensable role in this round of the AI revolution [6]. Brain dynamics modeling also urgently needs such a general-purpose programming framework.

To develop a general-purpose programming framework for brain dynamics modeling, we face several challenges.

- The first challenge comes from the complexity of the brain, which is organized modularly, hierarchically, and across multi scales [7]. This structural characteristic implies that researchers must build neural models at different levels (e.g., channel, neuron, network), and combine them freely across multiple levels for constructing larger models (e.g., neurons to networks, networks to functional systems). Current brain simulators typically focus on only one or two scales, e.g., spiking networks [8–12] or firing rate models [13–16]. Recently, NetPyNE [17, 18] and BMTK [19] have adopted descriptive languages to expand the modeling scales from channels to neurons and networks, but they still restrict users from defining models beyond predefined scales.
- The second challenge is the integration of different modeling needs. To elucidate brain functions comprehensively with computational models, we need to not only simulate neural activities, but also analyze the underlying mechanisms, and sometimes, we need to train models from data or tasks, implying that a general-purpose programming framework needs to be a platform to integrate multiple modeling demands. Current brain simulators mainly focus on simulation [20–22], and largely ignore training and analysis.
- The third challenge is achieving high running performance while maintaining programming convenience, which is particularly true for brain dynamics modeling, as its unique characteristics make it difficult to run efficiently within a convenient Python interface (see Results 2.4). The current popular approach for solving this challenge is code generation based on descriptive languages [22, 23]. However, this approach has intrinsic limitations regarding transparency, flexibility and extensibility [21, 22] (see Supplementary 1.1).
- The fourth challenge comes from the rapid development of the field. Brain dynamics modeling is relatively new and developing rapidly. New concepts, models, and mathematical approaches are constantly emerging, implying that a general-purpose programming framework needs to be extensible to take up new advances in the field conveniently.

In this paper, we propose BrainPy (“Brain Dynamics Programming in Python”, Fig. 1) as a solution to address all the above challenges. BrainPy provides infrastructure tailored for brain dynamics programming, including mathematical operators, differential equation solvers, universal model building interface, and object-oriented just-in-time (JIT) compilation based on the mature JIT platforms JAX [24], XLA [25] and Numba [26] (see Supplementary 1.2). Such infrastructure provides the flexibility for users to define brain dynamics models freely and lays foundations for BrainPy to build an integrative framework for brain dynamics modeling. First, BrainPy introduces a `DynamicalSystem` interface to unify diverse brain dynamics models. Models at any level of resolution can be defined as `DynamicalSystem` classes, which further can be hierarchically composed to create higher-level models. Second, BrainPy builds an integrated platform for studying brain dynamics models, where the same BrainPy model can be used for simulation, training (e.g., offline learning, online learning, or backpropagation training), and analysis (e.g., low-dimensional bifurcation analysis or high-dimensional slow point analysis). Third, through JIT compilation and dedicated operators, BrainPy achieves impressive performance for its code execution. The same models can be deployed into different devices (such as CPU, GPU and TPU) without additional code modification. Fourth, BrainPy is highly extensible. New extensions can be easily implemented as plug-in modules. Even the low-level primitive operators in the kernel system can be extended in the user-level Python interface. BrainPy is implemented in a robust continuous integration pipeline and is equipped with an automatic documentation building environment (see Supplementary 2.1). It is open-sourced at https://github.com/PKU-NIP-Lab/BrainPy. Rich tutorials and extensive examples are available at https://brainpy.readthedocs.io and https://brainpy-examples.readthedocs.io, respectively.

**Fig. 1:**
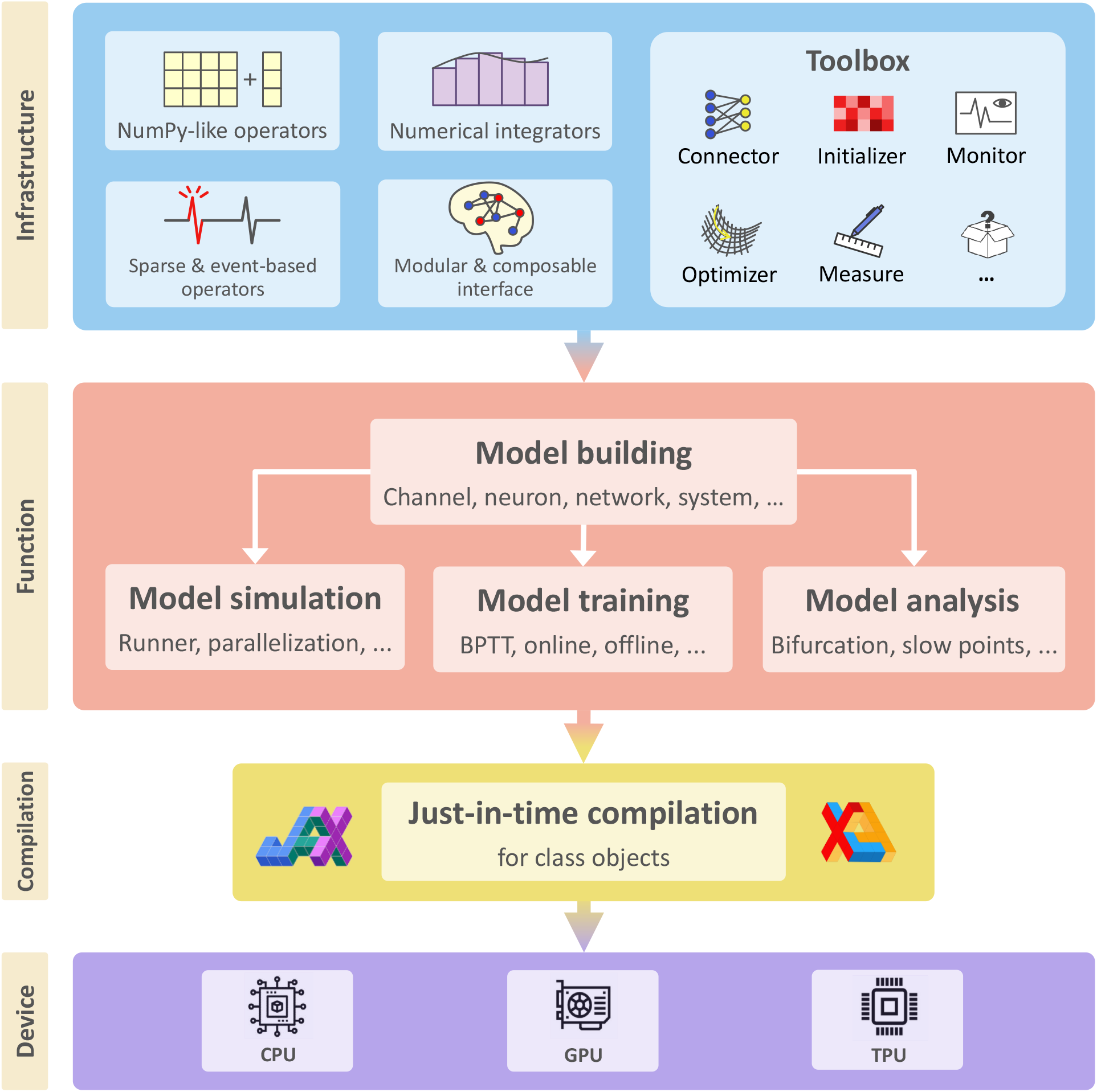
BrainPy is an integrative framework targeting general-purpose brain dynamics programming. **Infrastructure** panel. BrainPy provides infrastructure tailored for brain dynamics programming, including NumPy-like operators for computations based on dense matrices, sparse and event-based operators for event-driven computations, numerical integrators for solving differential equations, the modular and composable programming interface for brain dynamics model building, and a toolbox useful for brain dynamics modeling. **Function** panel. BrainPy provides an integrated platform for studying brain dynamics, including model building, simulation, training, and analysis. Models defined in BrainPy can be used for simulation, training, and analysis jointly. **Compilation** panel. Based on JAX [24] and XLA [25], BrainPy provides JIT compilation for Python class objects. All models defined in BrainPy can be JIT compiled into machine codes to achieve high-running performance. **Device** panel. The same BrainPy model can run on different devices including CPU, GPU, or TPU, without additional code modification.

## 2 Results

### 2.1 Infrastructure tailored for brain dynamics programming

To support its goal of becoming a general-purpose programming framework, BrainPy provides the infrastructure essential for brain dynamics modeling (see Fig. 1 “Infrastcture” panel). This infrastructure is a collection of inter-connected utilities designed to provide foundational services that enable users to easily, flexibly and efficiently perform various types of modeling for brain dynamics. Specifically, BrainPy implements: 1) mathematical operators for the conventional computations based on dense matrices and the event-driven computations based on sparse connections; 2) numerical integrators for various differential equations, the backbone of dynamical neural models; 3) a universal model building interface for constructing multi-scale brain dynamics models, and the associated JIT compilation for efficient running of these models; and 4) a toolbox specialized for brain dynamics modeling. They are outlined below.

First, BrainPy provides rich NumPy-like numerical functions and dedicated primitive operators for mathematical operations in brain dynamics modeling. Due to its simple and expressive syntax, successors of numerical libraries in Python prefer to follow NumPy [27]. The JAX framework has already provided an essential subset of NumPy functions capable of performing JIT transformations (see Supplementary 1.2), however, it still has many other functions, such as in-place updating and random sampling, which are significantly different from NumPy. To reduce the cost of learning a new set of computing languages, the BrainPy module `brainpy.math` supplements the missing or inconsistent NumPy numerical functions in JAX, including those for multidimensional arrays, mathematical operations, and random number generations (see Methods 4.1). NumPy-like functions effectively meet the computational needs involving dense matrices in brain dynamics modeling. However, brain dynamics models also feature event-based computation and sparse connections, which are unsuitable for direct implementation by NumPy-like operators. Therefore, BrainPy provides dedicated operators optimized for sparse and event-based computations in brain dynamics modeling (see Methods 4.1).

Second, BrainPy offers a repertoire of numerical solvers for solving differential equations. Differential equations are involved in most brain dynamics models. BrainPy’s numerical integration of differential equations is designed as a Python decorator. Users define differential equations as Python functions, whose numerical integration is accomplished by calling integrator functions, e.g., `brainpy.odeint()` for ordinary differential equations (ODEs), `brainpy.sdeint()` for stochastic differential equations (SDEs), and `brainpy.fdeint()` for fractional differential equations (FDEs). These integrator functions are designed to be general (see Methods 4.2), and most numerical solvers for ODEs and SDEs are provided, such as explicit Runge-Kutta methods, adaptive Runge-Kutta methods, and Exponential methods. For SDEs, BrainPy supports different stochastic integrals (Itô or Stratonovich) and different types of Wiener processes (scalar or multi-dimensional). As delays are ubiquitous in brain dynamics, BrainPy supports the numerical integration of delayed ODEs, SDEs, and FDEs with various delay forms (see Methods 4.2).

Third, BrainPy supports modular and composable programming and the associated object-oriented transformations. To capture the fundamental characteristics of brain dynamics, which are modular, multi-scaled, and hierarchical [7], BrainPy follows the philosophy that “any dynamical model is just a Python class, and high-level models can be recursively composed by low-level ones” (details will be illustrated in Results 2.2). However, such a modular and composable interface is not directly compatible with JIT compilers such as JAX and Numba, because they are designed to work with pure functions (see Supplementary 1.1). By providing object-oriented transformations (see Methods 4.3), including the JIT compilation for class objects and the automatic differentiation for class variables, models defined with the above modular and composable interface can also benefit from the powerful compilation supports of advanced JIT compilers.

Fourth, BrainPy offers a toolbox specialized for brain dynamics modeling (see Fig. 1 “Toolbox”). A typical modeling experiment involves multiple stages or processes, such as creating synaptic connectivity, initializing connection weights, presenting stimulus inputs, and analyzing simulated results. For the convenience of running these operations repeatedly, BrainPy presets a set of utility functions, including synaptic connection, weight initialization, input construction, and data analysis. However, this presetting does not prevent users from defining their utility functions in the toolbox.

### 2.2 Modular and composable programming for model building

Brain dynamics models have the key characteristics of being modular, multiscaled and hierarchical, and BrainPy designs a modular and composable programming paradigm to match these features. The paradigm is realized by the universal model building interface `DynamicalSystem`, which has the following appealing features.

First, the `DynamicalSystem` interface supports the definition of brain dynamics models at any organization level. Given a dynamical system, regardless of its complexity, users can implement it as a `DynamicalSystem` class. As an example, Fig. 2 b-1 demonstrates how to define a potassium channel model with Dyn≡icaisystem, in which the initialization function `__init__()` defines parameters and states, and the update function `update()` specifies how the states evolve. In this process, BrainPy toolbox can help users quickly initialize model variables, synaptic connections, weights and delays, and BrainPy operators and integrators can support users to freely define model updating logic. In a similar fashion, other dynamical models, such as neurons (e.g., discontinuous leaky integrate-and-fire (LIF) model [28], continuous FitzHugh–Nagumo model [29]), population models (e.g., Wilson-Cowan model [30], and ALN model [31]) and networks (e.g., continuous attractor neural network [32]), can be implemented by subclassing `DynamicalSystem` as standalone modules. For complex dynamical models, such as Hodgkin–Huxley (HH) type neuron models and large-scale cortical networks, they can either be defined as `DynamicalSystem` classes directly, or be composed by using sub-components (see below).

**Fig. 2:**
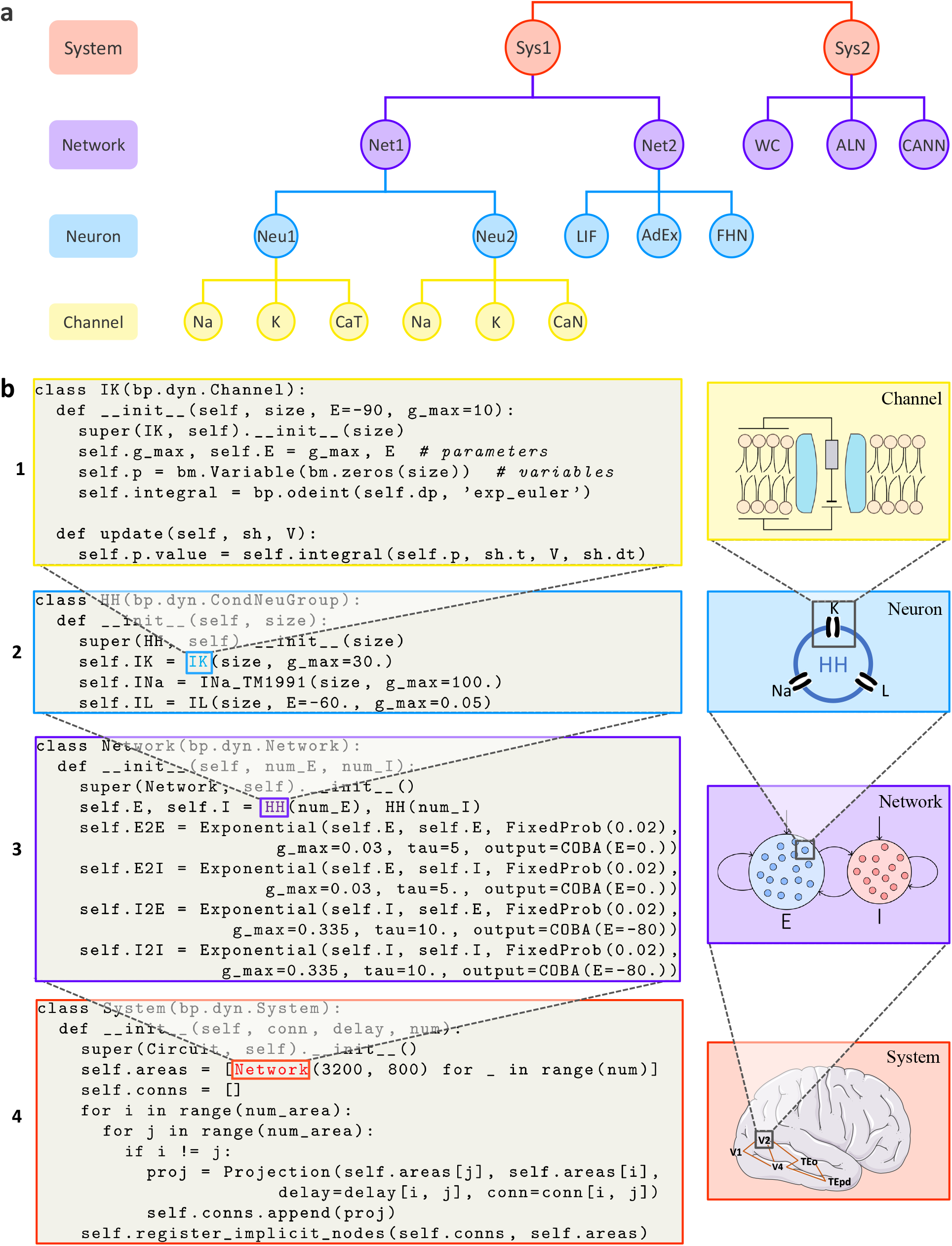
BrainPy supports modular and composable programming for building brain dynamics models. **a**. Illustrating the modular and hierarchical structure of brain dynamics models. A system-level model is composed of several network models (spiking or rate models). A network model is made up of many neuron models (HH or reduced neuron models). A neuron model is constructed from many ion channel models. **b**. An example demonstrating the construction of a hierarchical brain dynamics model using the interface `DynamicalSystem`. **b-1**, Defining an ion channel model as a subclass of `DynamicalSystem`, in which the function `__init__()` specifies the parameters and states, and the function `update ()` the updating rule of states. **b-2**, Defining an HH-typed neuron model by composing several ion channel models as a subclass of `DynamicalSystem`. **b-3**, Defining an E/I balanced network model by composing two neuron populations and their connections as a subclass of `DynamicalSystem`. **b-4**, Defining a ventral visual system model by composing several networks `V1, V2, V4, TEo`, and `TEpd` as a subclass of `DynamicalSystem`.

Second, the `DynamicalSystem` interface supports the definition of complex dynamical models in a composable manner. All models defined with `DynamicalSystem` can be used as modules to form more complicated high-level models. As an example, Fig. 2 b-2 demonstrates how an HH-type neuron model is created by combining multiple ion channel models (also see illustrations of “Neu1” and “Neu2” in Fig. 2a and Supplementary 3.1). Composable programming is the core of `DynamicalSystem`, and applies to almost all BrainPy models. For example, a synapse model consists of three components: synaptic dynamics (e.g., alpha, exponential, or dual exponential dynamics), synaptic output (e.g., conductance-based, current-based, or magnesium blocking-based), and synaptic plasticity (e.g., short-term or long-term plasticity). Composing different realizations of these components enables to create diverse kinds of synaptic models (see Supplementary 3.2). Similarly, various network models can be implemented by combining different neuron groups and their synaptic projections (see illustrations of “Net1” and “Net2” in Fig. 2a, a demonstration in Fig. 2 b-3, and more examples in Supplementary 3.3).

Third and remarkably, the `DynamicalSystem` interface achieves recursive composable programming [33], in term of that a model composed of lower-level components can recursively serve as a new component to form higher-level models. This property is highly useful for the construction of multi-scale brain models. Fig. 2b demonstrates an example of creating a complex model recursively: a ventral visual system model is created by interconnecting five E/I balanced network models (mimicking V1, V2, V4, TEo, TEpd brain areas, respectively; Fig. 2 b-4); each network model is formed by thousands of HH neurons (Fig. 2 b-3); and each HH neuron is further made up of multiple channel models (Fig. 2 b-2). It is worth pointing out that this recursive composition property is not shared by other brain simulators, for example, NetPyNE [18] and BMTK [19], which only support model definitions within the predefined scales. In contrast, BrainPy allows for flexible control of composition depth according to users’ needs. Moreover, for user convenience, BrainPy provides dozens of commonly used models, including channels, neurons, synapses, populations, and networks, as building blocks in the brainpy.dyn module to simplify the building of large-scale models (see Supplementary 3.4).

### 2.3 An integrative platform for model simulation, training, and analysis

Built upon the infrastructure, BrainPy provides an integrative platform, on which brain dynamics models can be simulated, trained, and analyzed jointly.

#### 2.3.1 Model simulation

BrainPy designs the interface `DSRunner` to simulate the dynamics of brain models (Method 4.4). `DSRunner` can be used to simulate models at any level, including channel (Fig. 3 a-1), neuron (Fig. 3 a-2), network (Fig. 3 a-3), and system (Fig. 3 a-4) levels.

**Fig. 3:**
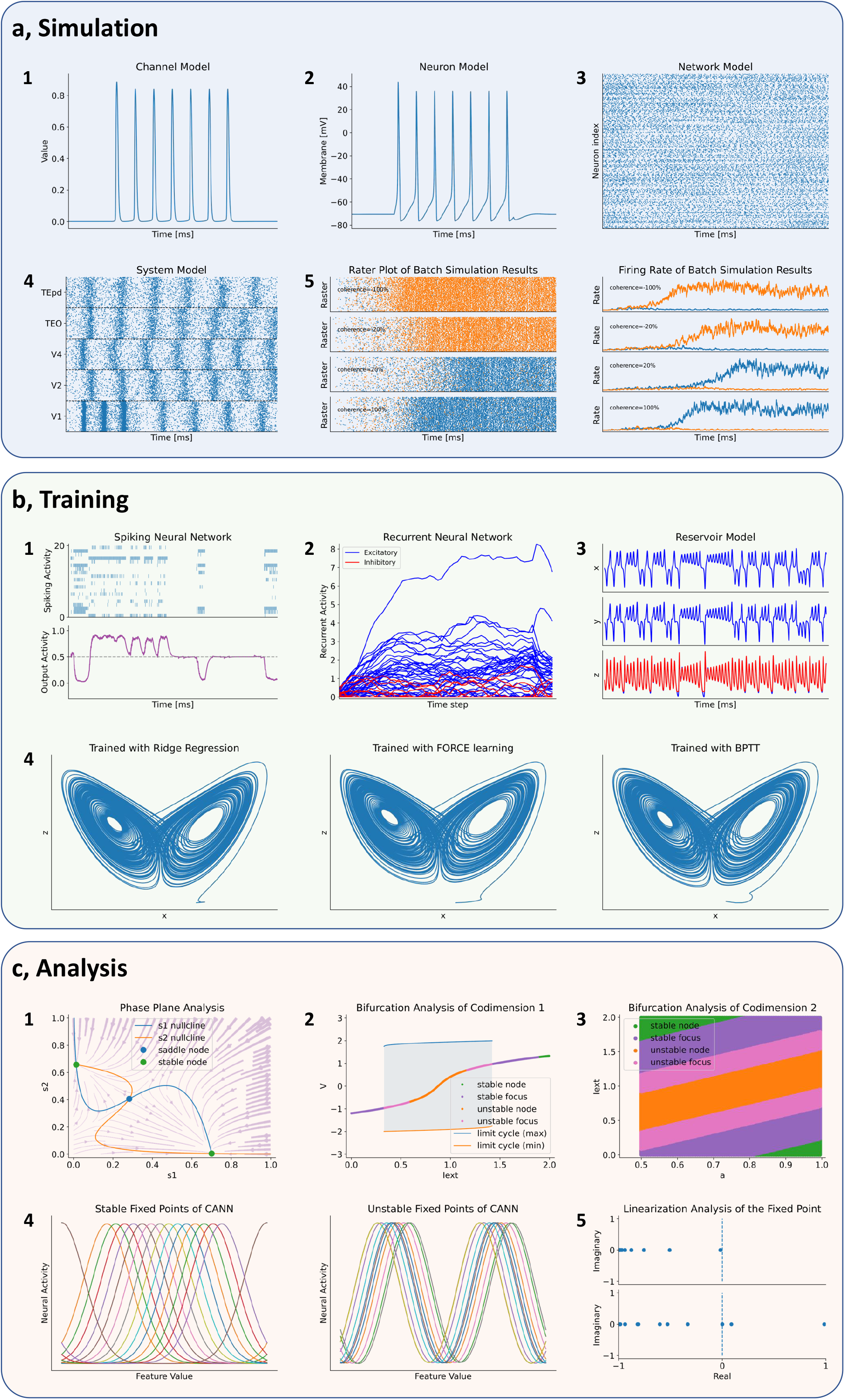
BrainPy provides an integrated platform for model simulation, model training, and dynamics analysis. **a**. The interface `DSRunner` supports the simulation of brain dynamics models at various levels. **a-1**, The simulation of the Potassium channel in Fig. 2 b-1. **a-2**, The simulation of the HH neuron model in Fig. 2 b-2. **a-3**, The simulation of the E/I balanced network, COBAHH [20] in Fig. 2 b-3. **a-4**, The simulation of the ventral visual system model in Fig. 2 b-4. **a-5**, Batch simulations of a spiking decision-making model with inputs of different coherence levels [34]. **b**. BrainPy supports the training of brain dynamics models from data or tasks. **b-1**, Training a spiking neural network [42] on an evidence accumulation task [64] using the backpropagation algorithm with `BPTT`. **b-2**, Training an artificial recurrent neural network model [65] on a perceptual decisionmaking task [66] with `BPTT`. **b-3**, Training a reservoir computing model [41] to infer the Lorenz dynamics with the ridge regression algorithm implemented in `OffiineTrainer`. *x,y* and *z* are variables in the Lorenz system. **b-4**, The classical echo state machine [67] is trained with multiple algorithms in BrainPy to predict the chaotic Lorenz dynamics, including ridge regression, FORCE learning, and BPTT. **c**. BrainPy supports automatic dynamics analysis for low- and high-dimensional systems. **c-1**, Phase plane analysis of a rate-based decision-making model [45]. **c-2**, Bifurcation analysis of codimension 1 of the FitzHugh–Nagumo model [29], in which the bifurcation parameter is the external input `Iext`. **c-3**, Bifurcation analysis of codimension 2 of the FitzHugh–Nagumo model [29], in which two bifurcation parameters `Iext` and a are continuously varying. **c-4**, Finding stable and unstable fixed points of a high-dimensional CANN model [32]. **c-5**, Linearization analysis of the high-dimensional CANN model [32] around one stable and one unstable fixed point.

In the simulation of brain dynamics models, it is often necessary to search intensively for parameter values to fit the experimental data, which is highly time-consuming. To speed up this process, BrainPy supports multiple ways to perform parallel simulations at different levels (see Supplementary 3.5), including parallelization based on multiple threads or microprocessors on a single machine, and parallelization across multiple devices. These parallelizations can also be used in a nested manner, for instance, by running a set of BrainPy models concurrently on different processors of multiple devices. Fig. 3 a-5 shows an example that demonstrates that a decision-making network model [34] with different external inputs can be simulated in parallel in BrainPy.

#### 2.3.2 Model training

The use of machine learning methods to train neural models is becoming a new trend and useful for studying brain functions [35–38]. BrainPy provides the `DSTrainer` interface to support this utility (Method 4.5). Different subclasses of `DSTrainer` provide different training algorithms, which can be used to train different types of models. For instance, the trainer `BPTT` implements the algorithm of backpropagation through time, which is helpful for training spiking neural networks (Fig. 3 b-1) and recurrent neural networks (Fig. 3 b- 2). Similarly, `offlineTrainer` implements offline learning algorithms such as ridge regression [39], and `onlineTrainer` implements online learning algorithms such as FORCE learning [40], and they are useful for training reservoir computing models (Fig. 3 b-3). In a typical training task, one may try different algorithms that can be used to train a model. The unified syntax for defining and training models in BrainPy enables users to train the same model using multiple algorithms. Fig. 3 b-4 demonstrates that a reservoir network model can be trained with three different algorithms: online, offline, and backpropagation, to accomplish a classical task of chaotic time series prediction [41].

Since the training algorithms for brain dynamics models have not been standardized in the field, BrainPy provides interfaces to support flexible customization of training algorithms (see Supplementary 3.6). Specifically, `offlineTrainer` and `onlineTrainer` provide general interfaces for offline and online learning algorithms respectively, and users can easily select the appropriate method by specifying the `fit_method` parameter in `OfflineTrainer` or `OnlineTrainer`. Users can also customize their own offline or online fitting methods by subclass-ing `offlineAigorithm` or `onlineAlgorithm`, in which they specify how model parameters are changed according to inputs, predictions, or target data. Furthermore, the interface `BPTT` is designed to capture the latest advances in backpropagation algorithms. For instance, it supports eligibility propagation algorithm [42] and surrogate gradient learning [43] for training spiking neural networks.

#### 2.3.3 Model Analysis

Analyzing model dynamics is as essential as model simulation and training, because it helps unveil the underlying mechanism of model behaviors. Traditionally, model analysis has been programmed separately from model simulation. Here, BrainPy provides a unified platform on which the model’s simulation, training, and analysis can be performed jointly. Specifically, given a dynamical system, BrainPy provides the interface `DSAnalyzer` for automatic dynamic analysis (Methods 4.6). Different classes of `DSAnalyzer` implement different analytical methods.

First, BrainPy supports phase plane and bifurcation analyses for lowdimensional dynamical systems. The phase plane is a classical and powerful technique for the analysis of dynamical systems and has been widely used in brain dynamics studies, including neuron models (e.g., Izhikevich model [44]) and population rate models (e.g., Wilson-Cowan model [30]). Fig. 3 c-1 shows an example where many features of phase plane analysis, including nullcline, vector field, fixed points, and their stability, for a complex rate-based decisionmaking model [45] are automatically evaluated by several lines of BrainPy code (see SI Listing 14). Bifurcation analysis is another utility of BrainPy, which allows users to easily investigate the changing behaviors of a dynamical system when parameters are continuously varying. Fig. 3 c-2 shows an example that demonstrates the stability changes of the classical FitzHugh–Nagumo model [29] with one parameter varying can be easily inspected by the bifurcation analysis interface provided in BrainPy. Similarly, bifurcation analysis of codimension-2 (with two parameters changing simultaneously; see Fig. 3 c-3 and SI Listing 16) can be performed with the same interface. BrainPy also supports bifurcation analysis for three-dimensional fast-slow systems, for example, a bursting neuron model [46]. This set of low-dimensional analyzers is performed numerically so that they are not restricted to equations with smooth functions, but are equally applicable to ones with strong and complex nonlinearity.

Second, BrainPy supports slow point computation and linearization analysis for high-dimensional dynamical systems. With powerful numerical optimization methods, one can find fixed or slow points of a high-dimensional nonlinear system [36, 38, 47, 48]. By integrating numerical methods such as gradient descent and nonlinear optimization algorithms, BrainPy provides the interface `slowPointFinder` as a fundamental tool for high-dimensional analysis. To illustrate this, we applied `slowPointFinder` to a CANN network model [32] and found that it can successfully find a line of stable and unstable attractors in CANN (see Fig. 3 c-4). Furthermore, the linearized dynamics around fixed points can be easily inspected and visualized with a single line of functional call in BrainPy (see Fig. 3 c-5 and SI Listing 17).

### 2.4 Efficient performance of BrainPy

Simulating dynamical models efficiently in Python is challenging [9, 22]. To resolve this problem, BrainPy leverages the JIT compilation of JAX/XLA and exploits dedicated primitive operators to accelerate the model running.

In contrast to deep neural networks (DNNs), which mainly consist of computation-intensive operations (such as convolution and matrix multiplication), brain dynamics models are usually dominated by memory-intensive operations. Taking the classical LIF neuron model [28] as an example, its computation mainly relies on operators such as addition, multiplication and division. Fig. 4a compares the running times between a LIF model and a matrix multiplication under the floating point operations (FLOPs), which shows that although both modes have the same theoretical complexity, the LIF model runs 10 times longer on both CPU and GPU than the matrix multiplication (Fig. 4a and SI Fig. 4), thus demonstrating that large overheads exist when running brain dynamics models in Python. Particularly, these overheads become overwhelming when simulating large-scale brain networks as they increase rapidly with the number of operators in the model. In BrainPy, we exploit the JIT compilation to reduce these overhead costs dramatically. The JIT compilation lowers the dynamic Python code into the static machine code during runtime, which can significantly reduce the time cost of Python interpretation. In particular, BrainPy takes advantage of JAX, which achieves JIT compilation based on XLA (see Supplementary 1.2). The XLA JIT engine has optimizations specialized for memory-intensive operators, such as operator fusion, making it highly suitable for processing brain dynamics models. Fig. 4a shows that by leveraging the JIT compilation, the LIF model has a running speed comparable to that of the matrix multiplicative operation `dot` on the CPU and achieves an even better performance on the GPU (SI Fig. 4). We further demonstrate that for a large-size E/I balanced network model COBA [20], the JIT compilation accelerates the running speed 10 times on both the CPU and GPU compared to no JIT compilation (see Fig. 4b and SI Fig. 5).

**Fig. 4:**
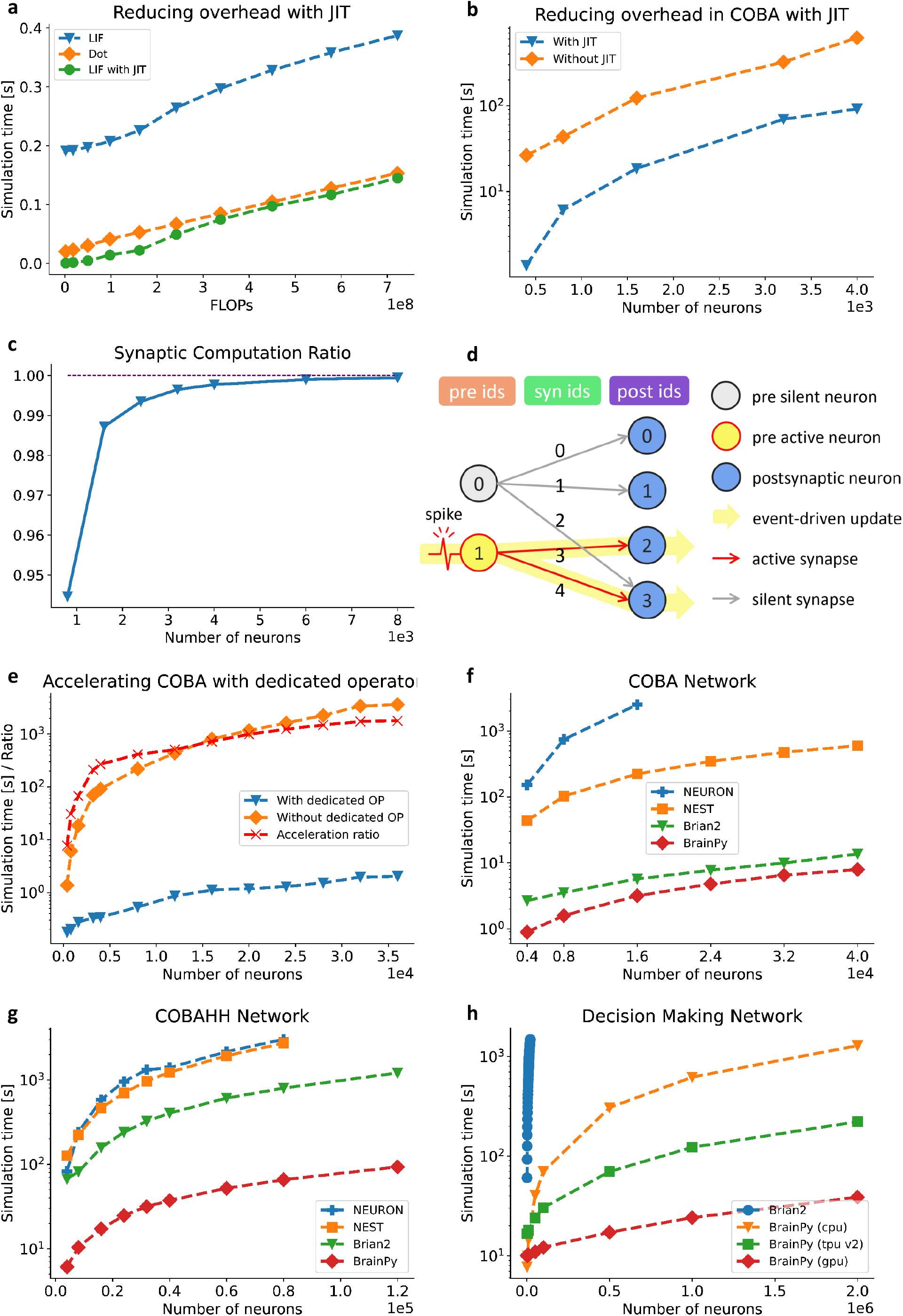
BrainPy achieves efficient performance through JIT compilation and dedicated primitive operators. **a**, Performance comparison between a LIF neuron model [28] and a matrix multiplication *Wx* (*W* ∈ ℝ^*m*×*m*^ and *x* ∈ ℝ^*m*^). By adjusting the number of LIF neurons in a network and the dimension *m* in the matrix multiplication, we compare two models under the same FLOPs. Without JIT compilation, the LIF model shows ten times slower than the matrix multiplication. After lowering the whole LIF network into the CPU device through JIT compilation, two models show comparable running speeds. **b**, The JIT compilation accelerates a classical E/I balanced network COBA [20] ten times on CPU. **c**, Without dedicated operators, more than 94% of the time is spent on synaptic computations when simulating a COBA network model. The ratio increases with the network size. **d**, Illustration of data conversion in a typical sparse synaptic computation. The state of a post-synaptic neuron or the synapse strength is updated based on the presynaptic spikes, corresponding to the “pre-to-post” or “pre-to-syn” event-driven computation. Similarly, synapse-level events can drive the computation of post-synaptic variables, i.e., the “syn-to-post” event-driven computation. **e**, The COBA network model with event-based operators is accelerated by up to three orders of magnitude compared to that without dedicated operators. **f**, Speed comparison of NEURON, NEST, Brian2, and BrainPy on the benchmark model COBA. **g**, Speed comparison of NEURON, NEST, Brain2, and BrainPy on the benchmark model COBAHH. **h**, Speed comparison of a spiking decision-making network [34] on CPU, GPU or TPU. In the experiments in **a**, differential equations were solved using the Euler method with just one time step. For other experiments, dynamical equations were solved using the Exponential Euler method with a step size of 0.1 ms. All simulations were run on Python 3.8 installed on Ubuntu 20.04.2 LTS. Simulations run on CPU were carried out with Intel(R) Core(TM) i7-6700K CPU @ 4.00 GHz. GPU benchmarking is based on NVIDIA Tesla V100. Google Colab free TPU v2 is used for TPU benchmarking. The versions of NEURON, NEST, and Brian2 used are 8.2.0, 2.20.0, and 2.5.1, respectively.

Other essential features distinguishing brain dynamics models from DNNs are that they usually have sparse connections and perform event-driven computations. For instance, neurons in a network are typically connected with each other with a probability smaller than 0.2 [49], and the state of a post-synaptic neuron is updated only when the event of presynaptic spiking arrives. These unique features greatly hinder the efficiency of brain model simulation by using conventional NumPy-like operators, even with the aid of JIT compilation. Fig. 4c shows that when implementing a COBA network model with NumPy-like operators, more than 94% of the simulation time is spent on synaptic computations, which worsens as the network size increases. To solve this problem, BrainPy designs dedicated primitive operators to accelerate the event-based computations embedded in sparsely connected networks. These dedicated operators include transformations among variables corresponding to presynaptic neurons (“pre”), post-synaptic neurons (“post”), and synapses (“syn”) (see Fig. 4d). In BrainPy, we predefine a dozen of operators for these “pre-to-syn”, “syn-to-post” and “pre-to-post” data transformations (Methods 4.1), which can significantly reduce the time consuming on synaptic computations. Fig. 4e illustrates that these dedicated event-based operators accelerate the classical COBA network thousands of times. Similar accelerations were achieved on the GPU (SI Fig. 6). By examining the proportion of time spent on synaptic computations, we found that with dedicated operators the time cost is only half that spent on the entire network simulation (SI Fig. 7), and this ratio no longer increases with the network size.

To formally assess the running efficiency of BrainPy, we compared it with other popular brain simulators, including Brian2, NEST, and NEURON (see Supplementary 2.2 for the selection of simulators for comparison). The benchmark models are the standard E/I balanced networks with LIF- or HH-typed neuron, i.e., COBA and COBAHH networks [20]. The comparison results are summarized in Fig. 4f and Fig. 4g, which show that BrainPy is highly efficient and achieves the best performances on both benchmark tasks, and the superiority of BrainPy increases as the network size increases.

Remarkably, BrainPy has a very appealing property not shared by other brain simulators, i.e., “code once, run everywhere”. This advantage comes from the fact that JAX and XLA are compatible with most operating systems, including macOS, Linux, and Windows, and their JIT compilations can automatically translate Python codes into efficient machine codes for CPU, GPU, or TPU devices, which means that the same model in BrainPy can run on different accelerators without additional code modification. In Fig. 4h, we run the same code for a decision-making network model [34] on the CPU, GPU, and TPU separately, and observe that the running speeds on the GPU and TPU are tens of times faster than that on the CPU. We also implemented this decision-making model with Brian2 on the CPU (although it was stated that Brian2 could target on GPU via code generation with GeNN [50, 51], we could not run this model on a GPU with Brian2 successfully), and observed that the running speed of Brian2 was significantly slower than that of BrainPy. In particular, the time cost of BrainPy increases slightly with network size, whereas the time cost of Brian2 increases dramatically.

### 2.5 Extensible architecture of BrainPy

Brain science, as well as brain dynamics modeling, is progressing rapidly. Along with the gigantic projects on brain research worldwide, new data and knowledge about brain structures and functions are constantly emerging, which impose new demands on brain dynamics modeling frequently, including, for instance, the simulation and analysis of large-size neural circuits, and the training of neural models based on recorded neural data. To be a general-purpose brain dynamics programming framework, the architecture of the framework must be extensible to conveniently take up new advances in the field. Current brain simulators based on descriptive languages have difficulty achieving this goal, since the extension of a new component through the descriptive interface needs to be done in both high- and low-level programming languages (see Supplementary 1.1). Through the elaborate architecture design, BrainPy enables easy extension with new infrastructure, new utility functions, and new machine learning methods, all performed in our convenient Python interface.

First, for infrastructure (see Fig. 1 “Infrastructure” panel), BrainPy provides a convenient way of customizing a new tool as defining a new subclass. For example, to add a new Runge-Kutta integrator, it can be done by subclassing `ExplicitRKIntegrator` and providing the Butcher tableau; to define a new connector, it only requires to inherit from `TwoEndConnector` and implement two functions: `__init__()` (initializing connection parameters) and `build_conn()` (specifying how the connections are established between two populations). To address the known difficulty of extending custom operators, BrainPy provides an interface that enables users to write and register an operator directly with Python syntax. Since models and modeling methods have not yet been standardized in the field, the abstraction and summarization of primitive operators for brain dynamics modeling are lacking. For this reason, although BrainPy provides dozens of dedicated operators, it would be too soon to establish a complete operator library for brain dynamics modeling. Therefore, BrainPy provides an `XLACustomOp` interface to simplify operator customization. Specifically, to customize a primitive operator, users need to subclass `XLACustomOp` and implement two Python functions: the abstract evaluation function `evai_shape()` and concrete computation function `con_compute()` (see Methods 4.7). Notably, this differs from the operator customization in the original JAX system, where low-level operators must be implemented through XLA custom call using C++ codes. We confirmed that operators customized through the BrainPy interface have comparable and even better performance than those written in C++ (see SI Fig. 8).

Second, for functional modules (see Fig. 1 “Function” panel), BrainPy enables an extension of a new module with BrainPy infrastructure, as the latter can be arbitrarily fused, chained, or combined to create new functions. For example, an analysis toolkit can be customized with NumPy-like and dedicated operators. Moreover, all customizations in BrainPy can benefit from the acceleration of JIT compilation, and users’ attention only needs to focus on the functions to be extended.

Third, for interactions with AI, BrainPy supports the easy extension of new machine learning methods. Machine learning approaches are becoming important tools for brain dynamics modeling [52]. Existing brain simulators have difficulty incorporating the latest advances in machine-learning research (see Supplementary 1.1). Built on top of the machine learning system JAX, BrainPy has the advantage of being linked to the latest developments in machine learning. JAX has a rich ecosystem of machine learning, including deep neural networks [53–55], graph neural networks [56], reinforcement learning [57], and probabilistic programming [58]. To integrate this rich machine learning ecosystem as part of the users’ program, BrainPy is designed to be compatible with other JAX libraries. For example, the array structure `brainpy. math.jaxArray` in BrainPy is allowed for the bidirectional exchange of data in JAX without any data copying (see Supplementary 3.7), and the object-oriented transformations of BrainPy can be applied to pure functions, thus enabling most JAX libraries with a functional programming style can be directly used as a part of the BrainPy program (see examples in Supplementary 3.8).

## 3 Discussion

A long-standing concern in brain dynamics modeling is that we lack a general-purpose programming framework that can support users to freely define brain dynamics models across multiple scales, comprehensively perform simulation, optimization and analysis of the built models, and conveniently prototype new modeling methods. By utilizing the JIT compilation technology, we developed BrainPy as a candidate for such a general-purpose programming framework, which has the following appealing properties:

- **Pythonic programming**. In contrast to other brain simulators which only provide Pythonic interface [8–12, 17–19, 21–23], BrainPy enables Pythonic programming. It allows users to implement and control their models directly using native Python syntax, which implicating high transparency to users. This transparency is crucial for research, as standard Python debugging tools can be integrated into the implementation process of novel models, and is also appealing for education.
- **Easy-to-lean-and-use**. By prioritizing usability as the top priority, BrainPy provides NumPy-like operators and decorator-based differential solvers to alleviate the learning cost of mathematical operations, and offers a modular and composable interface to simplify the building of hierarchical brain dynamics models.
- **Integrative platform**. BrainPy allows unprecedentedly integrated studying of brain dynamics models, in term of that model simulation, training, and analysis can be done jointly.
- **Intrinsic flexibility**. Inspired by the success of general-purpose programming in Deep Learning [4, 5], BrainPy provides not only functional libraries but also infrastructure. This is essential for users to create models and modeling approaches beyond the predefined assumptions of existing libraries.
- **Efficient performance**. BrainPy models are JIT compiled into machine codes to achieve extremely fast running speeds, and the same code can run on different devices, including CPUs, GPUs, and TPUs.
- **Extensible architecture**. BrainPy has an extensible architecture. New primitive operators, utilities, functional modules, machine learning approaches, and others, can be easily customized through our Python interface.

Nevertheless, to fulfil all demands of brain dynamics modeling, there are future works to be done to complete the ecosystem of BrainPy. These include, for instance, to support the efficient implementation of multi-compartment neurons needed for biological detailed modeling [59], to develop parallel primitive operators and memory-efficient algorithms for ultra-large brain simulation (up to billions of neurons) [60], and to provide the JIT compilation for neuro-morphic devices for deploying BrainPy models onto neuromorphic computing systems [61]. We hope that BrainPy can contribute to the research in brain science and eventually serve as a general-purpose framework for brain dynamics modeling.

## 4 Methods

### 4.1 Mathematical operators for brain dynamics modeling

Brain dynamics modeling involves the computation based on dense matrices and the event-driven computation based on sparse connections. BrainPy provides mathematical operators for these two kinds of computations.

#### 4.1.1 Dense matrix operators

NumPy [27] has been a gold standard in Python numerical computing for manipulating and operating on dense matrices (they are called “arrays” in NumPy). Our dense matrix operators attempt to follow NumPy’s syntax. Specifically, we provided a data structure `brainpy.math.jaxArray` which is consistent with NumPy’s `ndarray` structure, and a series of mathematical operations for `JaxArray` which are similar to those for `ndarray`.

`jaxArray` is a wrapper of JAX’s `DeviceArray` data, but it supports essential features of NumPy’s `ndarray` structure, including attributes like the number of axes (`.ndim`) and the dimensions (`.shape`), and build-in functions such as comparison, summation, and in-place updating (see the comparison in SI Table 1).

Mathematical operators for `JaxArray`, such as indexing, slicing, sorting, rounding, arithmetic operations, linear algebraic functions, and Fourier transform routines, are all supported. Many of these operators (nearly 85 percent) are directly implemented through the NumPy-like functions in JAX, while BrainPy provides dozens of important NumPy APIs missing or inconsistent in JAX (see the comparison in SI Table 1).

Moreover, to unify random number generators, BrainPy implements most of the random sampling functions in NumPy, including its univariate distributions, multivariate distributions, standard distributions, and utility functions (see the comparison in SI Table 1).

#### 4.1.2 Event-driven operators

Currently, event-driven operators in BrainPy are involved in transformations among pre-synaptic neuronal data, synaptic data, and post-synaptic neuronal data (see illustrations in Fig. 4d). Specifically, BrainPy provides operators for mapping pre-synaptic neuronal data to its connected synaptic dimension (“pre-to-syn”), arithmetic operators including summation, product, max, min, mean, and softmax for transforming synaptic data to post-synaptic neuronal dimension (“syn-to-post”), and arithmetic operators such as summation, product, max, min, and mean for directly computing post-synaptic data from pre-synaptic events (“pre-to-post”). Moreover, BrainPy provides event-based matrix multiplication operators with sparse connections.

### 4.2 Numerical solvers for differential equations

To meet the large demand for solving differential equations in brain dynamics modeling, BrainPy implements numerical solvers for various differential equations, including ordinary differential equations (ODEs), stochastic differential equations (SDEs), fractional differential equations (FDEs), and delay differential equations (DDEs). In general, numerical integration in BrainPy is just a Python decorator. Users define a differential equation as a Python function; then, the numerical integration of this Python function is implemented through a simple decorator, e.g., @brainpy.odeint for ODEs, @brainpy.sdeint for SDEs, and @brainpy.fdeint for FDEs (see the following text). Moreover, to be compatible with models that have discontinuous dynamics, the numerical solver in BrainPy returns a function which performs one integration step, that is, the numerical integration function defines the system evolution of *X*(*t*) → *X*(*t* + *dt*), where *X* is the state, *t* the current time, and *dt* the integration step. Thus, the discontinuous part of the model can be updated in time after performing one integration step.

#### 4.2.1 ODE numerical solvers

For a two-dimensional ODE system,

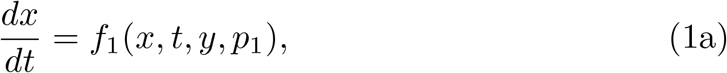

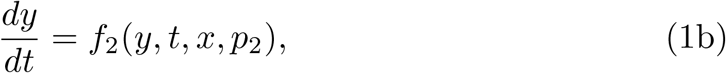

we can define it as a function as

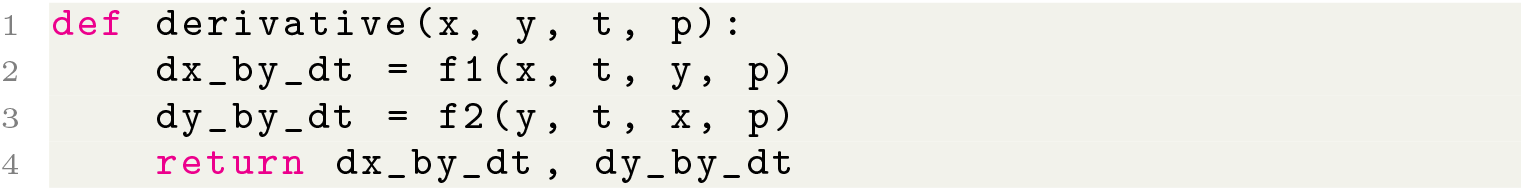

where `t` denotes the time variable, `p1` and `p2` after `t` denote the parameters needed in this system, and `x` and `y` before `t` represent the dynamical variables. In the function body, the derivative of each variable can be customized according to the user’s need (`f1` and `f2`). After defining the derivative function, numerical integration can be implemented using a simple decorator @brainpy.odeint(method, dt),

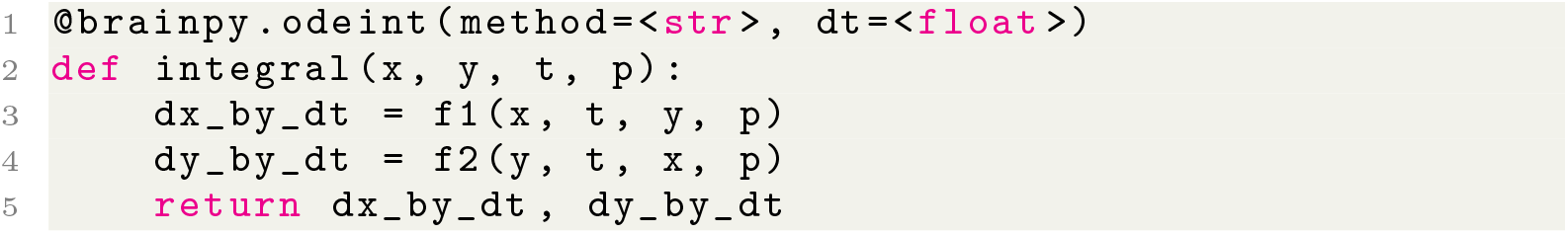

where `method` denotes the numerical method used to integrate the ODE function (use the “keyword” listing in SI Table 2 to specify `method`), and `dt` controls the initial or default numerical integration step.

A variety of numerical integration methods for ODEs, including Runge-Kutta, adaptive Runge-Kutta, and Exponential methods, are supported in BrainPy (see SI Table 2).

#### 4.2.2 SDE numerical solvers

Same as the ODE system, any SDE system expressed as

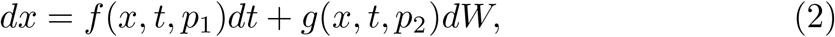

can be defined as two Python functions:

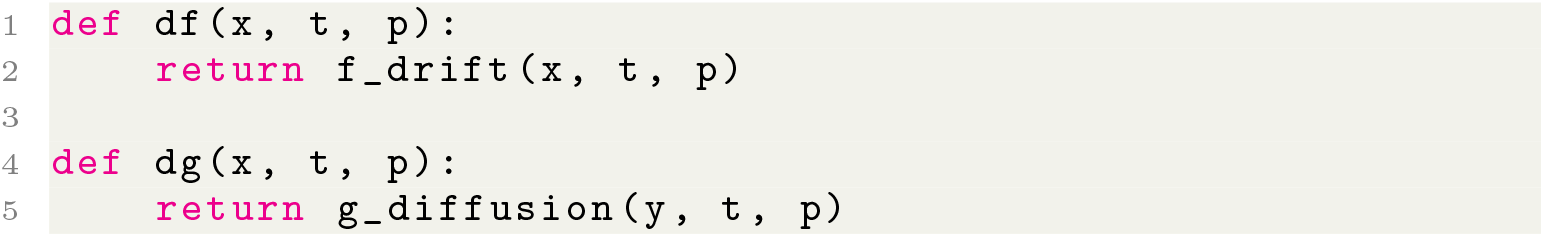

Specifically, the drift and diffusion coefficients in the SDE system are defined in the `f_drift` and `g_diffusion` functions. For the SDE function with scalar noise, the size of the return data *df* and *dg* should be the same, for example, *df* ∈ *R^d^* and *dg* ∈ *R^d^*. However, for a more general SDE system, it usually has multi-dimensional driving Wiener processes:

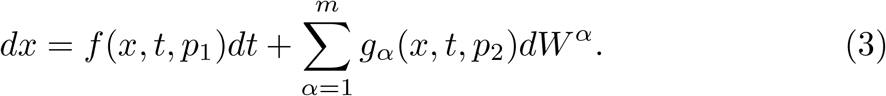

For such an m-dimensional noise system, the coding schema is the same as the scalar one, except that the return size of *dg* is multi-dimensional, for example, *df* ∈ *R^d^* and *dg* ∈ *R*^*d×m*^. SDEs have two ways of integral: Itô and Stratonovich stochastic integrals [62], and BrainPy supports both of them. Specifically, the numerical integration of SDEs is performed by `brainpy.sdeint`(`f, g, method, dt, wiener_type, integral_type`):

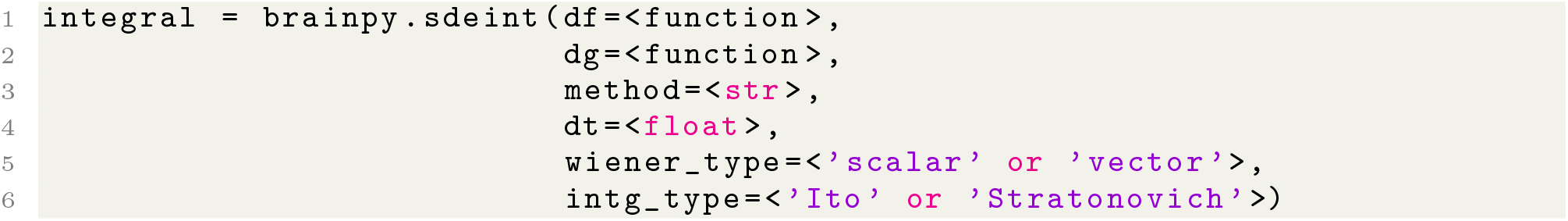

where `method` specifies the numerical solver (can be chosen by the “keyword” listing in SI Table 3), `dt` the default integral step, `wiener_type` the type of Wiener process (SCALAR_WIENER or `VECTOR_WIENER`), and `integral_type` the integral type (`ITO_SDE` or `STRA_SDE`).

#### 4.2.3 FDE numerical solvers

The numerical integration of FDEs is very similar to that of ODEs, except that the initial value, memory length, and fractional order should be provided. Given the fractional-order differential equation

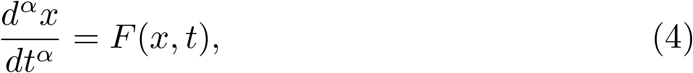

where the fractional order 0 < *α* ≤ 1. BrainPy supports its numerical integration with the following format of

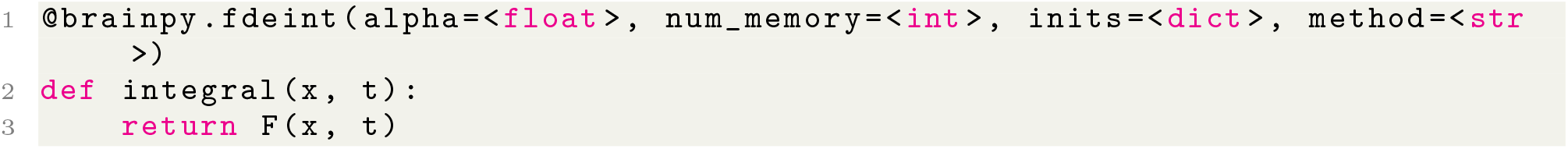

BrainPy supports two types of FDE, that is, the Caputo derivative and Grünwald-Letnikov derivative (see SI Table 4). Caputo derivatives are widely used in neuroscience modeling [63]. However, the numerical integration of Caputo derivatives has a high memory consumption, because it requires to integrate over all past activities. This implies that FDEs with the Caputo derivative definition cannot be used to simulate realistic neural systems. However, the numerical method for Grünwald-Letnikov FDEs, `brainpy.fde. GLshortMemory`, is highly efficient, because it does not require an infinite memory length for numerical solutions. Because of the decay property of the weighted coefficients for past activities, `brainpy.fde.GLshortMemory` implements a short memory length to reduce computational time. With the increasing length of short memory, the accuracy of the numerical approximation will increase. Therefore, `brainpy.fde.GLShortMemory` can be applied to real-world neural modeling.

#### 4.2.4 DDE numerical solvers

Delays occur in any type of differential equations. In a realistic neural modeling, delays are often inevitable. BrainPy supports equations with variables of constant delays, like

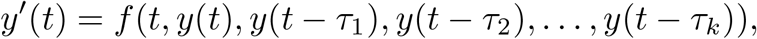

where the time lags *τ_j_* are the positive constants. It also supports systems with state-dependent delays, e.g.,

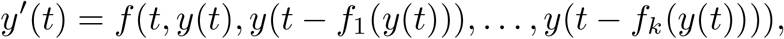

where *f_k_* is the function that computes the delay length by the system state *y*(*t*). For neutral typed equations in which delays appear in derivative terms,

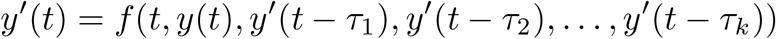

BrainPy also supports its numerical integration.

BrainPy, in particular, implements interfaces to define these various delay variables. `brainpy.math.TimeDelay` and `brainpy.math.LengthDelay` are provided to support state-dependent variables, and `brainpy.math.NeuTimeDelay` and `brainpy.math.NeuLenDelay` are implemented to model neutral delay variables. Moreover, the differential equations with delayed variables are intrinsically supported in each integral functions. For delay ODEs, users can use `brainpy.odeint`(`state_delays, neutral_delays`, …) with the specification of `state_delays` for state-dependent delay variables and `neutral_delays` for neutral delay variables. Similarly, the numerical integration of delay SDEs and FDEs should utilize `brainpy.sdeint`(`state_delays`, …) and `brainpy.fdeint`(`state_delays`, …), respectively, with an explicit declaration of statedependent delay variables. Note that we currently do not support neutral delays for delay SDEs and FDEs. However, users can easily customize their supports for equations with neutral delays.

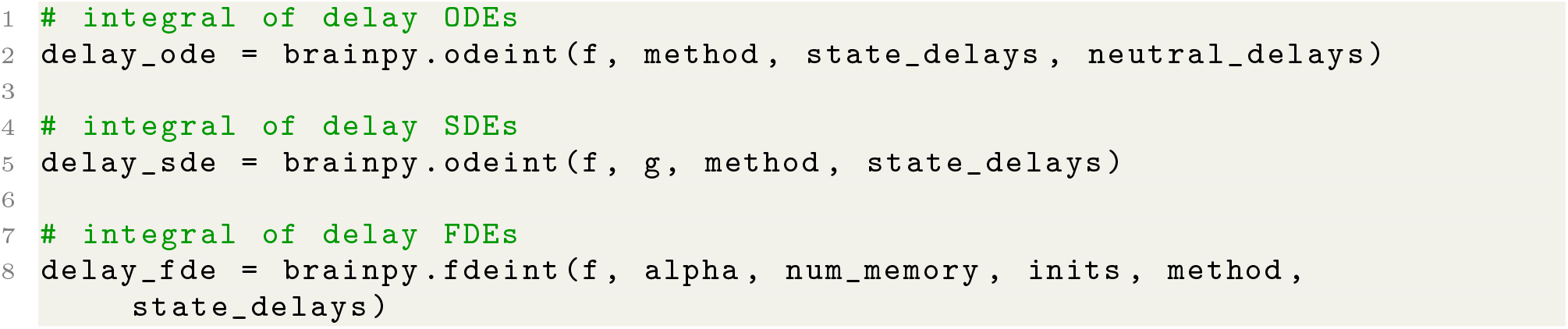

### 4.3 Object-oriented JIT compilation and automatic differentiation

Under minimal constraints and assumptions, BrainPy enables the JIT compilation for class objects. These assumptions include the following:

- The class for JIT must be a subclass of `brainpy.Base`.
- Dynamically changed variables in the class must be labeled as `brainpy.math.Variabie`. Otherwise, they will be compiled as constants and cannot be updated during the program execution.

To take advantage of the JIT compilation, we can directly apply `brainpy.math. jit()` onto the instantiated class objects, or functions of a class object.

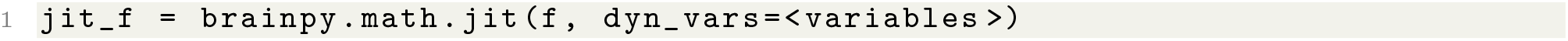

Automatic differentiation is fundamental in machine learning algorithms. BrainPy supports the automatic differentiation of class variables for its modular and composable programming paradigm, with a simple and elegant syntax.

`brainpy.math.grad()` takes a function/object (*f*: ℝ^*n*^ → ℝ, which returns a scalar value) as the input and returns a new function (*∂f* (*x*) → ℝ^*n*^) which computes the gradient of the original function/object.

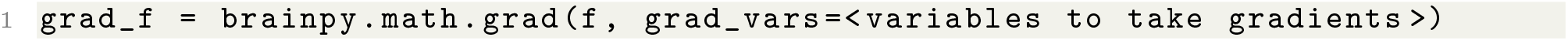

`brainpy.math.vector_grad()` takes vector-valued gradients for a function/object (*f*: ℝ^*n*^ → ℝ^*m*^). It returns a new function (*∂f* (*x*): ℝ^*m*^ → ℝ^*n*^) which evaluates the vector-Jacobian products.

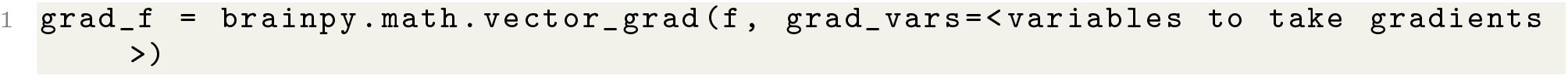

Another way to take gradients of a vector-output value is using `brainpy.math.jacobian()`. It aims to automatically compute the Jacobian matrices *•f* (*x*) ∈ ℝ^*m*×*n*^ by the given function/object *f*: ℝ^*n*^→ℝ^*m*^ at the given point of *x* ∈ ℝ^*n*^.

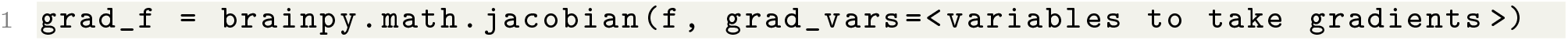

### 4.4 Simulation of brain dynamics models in BrainPy

BrainPy provides `brainpy.DSRunner` for simulating brain dynamics models. `DsRunner` has three functions: 1) running models over time, 2) presenting inputs for target variables during model running, 3) and monitoring history values of the interested variables.

The syntax of `brainpy.DSRunner` is simple. It receives the target model, the wanted inputs, and the interested monitors.

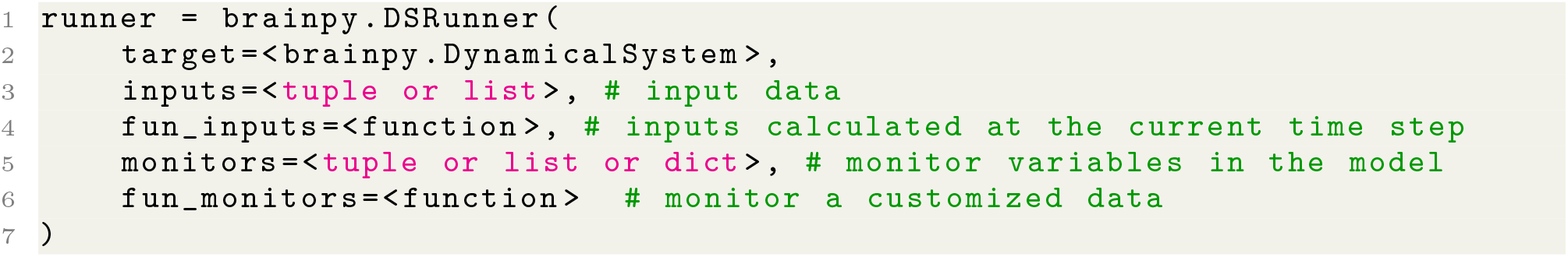

After initializing a runner, users can call `.run(duration)` function to perform a simulation according to the simulation duration the user specifies. `.run()` function can be repeatedly called for running a long time duration.

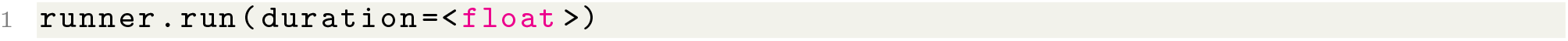

All history values of the monitored data are gathered in `runner.mon`.

### 4.5 Training of of brain dynamics models in BrainPy

Similar with model simulation, BrainPy provides `brainpy.train.DSTrainer` for training brain dynamics models. Different training methods are implemented in different `DSTrainer` subclasses. Currently, BrainPy provides three different training algorithms. They are the online training method which implemented in `brainpy.train.OnlineTrainer`, the offline training method which coded in `brainpy.train.OfflineTrainer`, and the backpropagation algorithm which provided in `brainpy.train.BPTT`.

Similar to `DSRunner`, initializing a `DSTrainer` needs to provide the target model, and it supports inputs and monitors too. For `OnlineTrainer` and `OfflineTrainer`, they receive a `fit_method` parameter for choosing fitting method. For example,

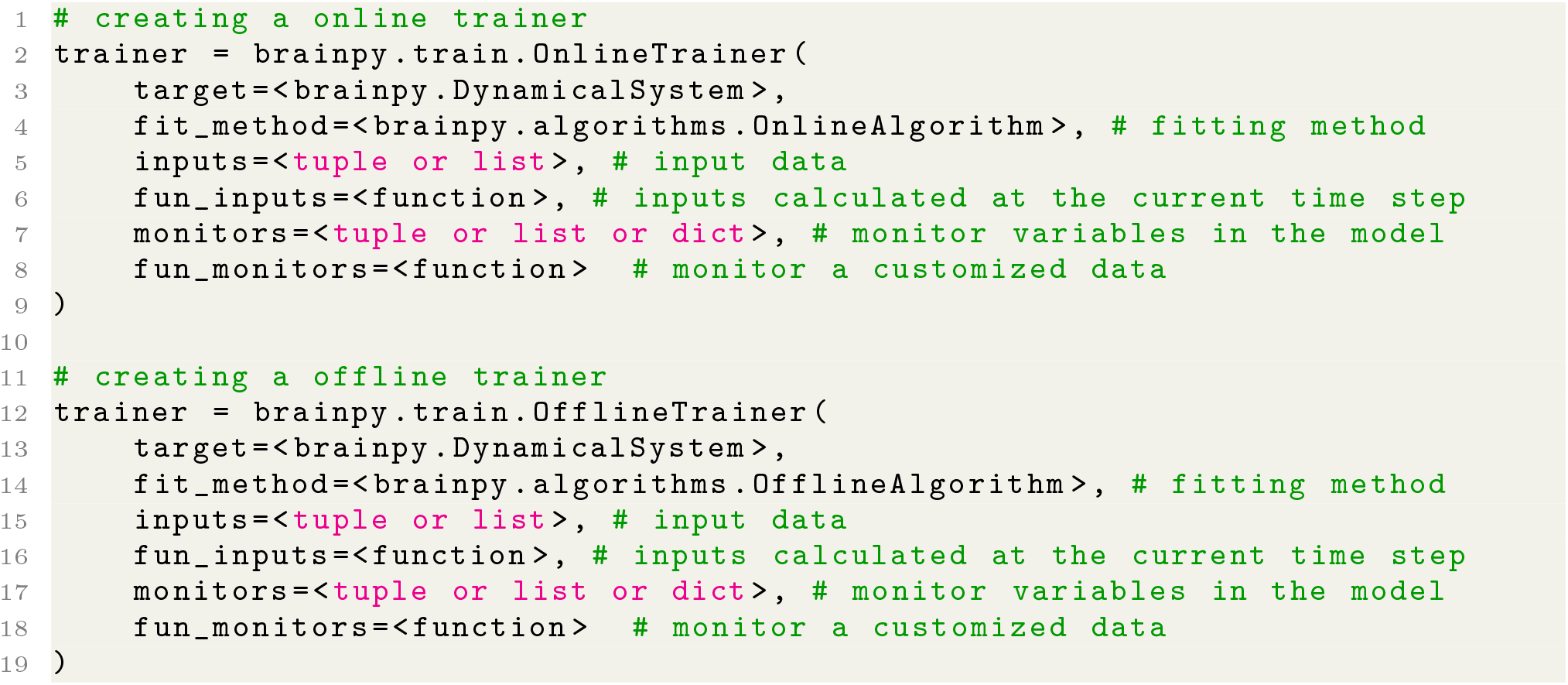

Different from `OnlineTrainer` and `offiineTrainer, BPTT` needs two more parameters, which are `loss_fun` for calculating training losses and `optimizer` for the network parameter optimization.

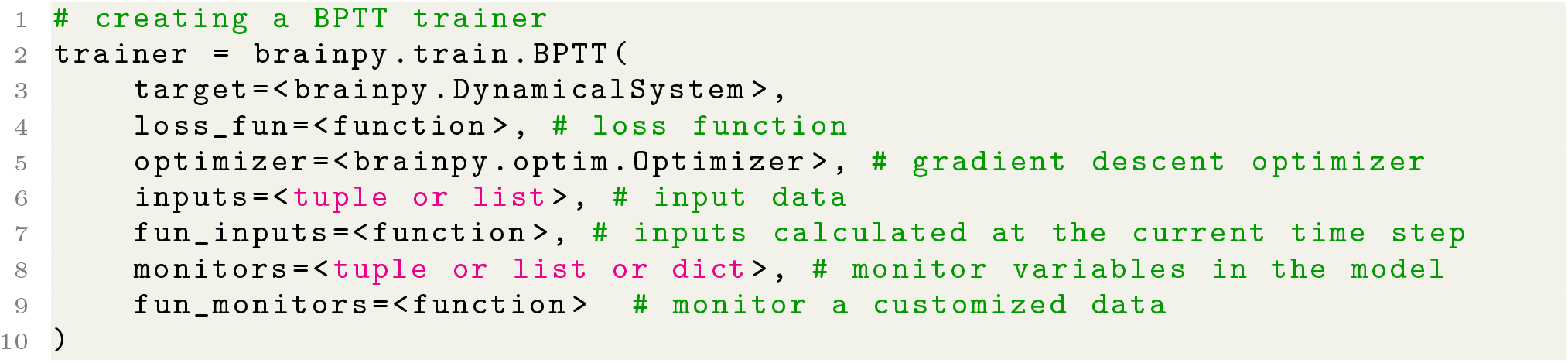

All trainers in BrainPy have two useful functions, one is the `.fit()` for fitting model with the data, the other is the `.predict()` for predicting outputs according to the inputs.

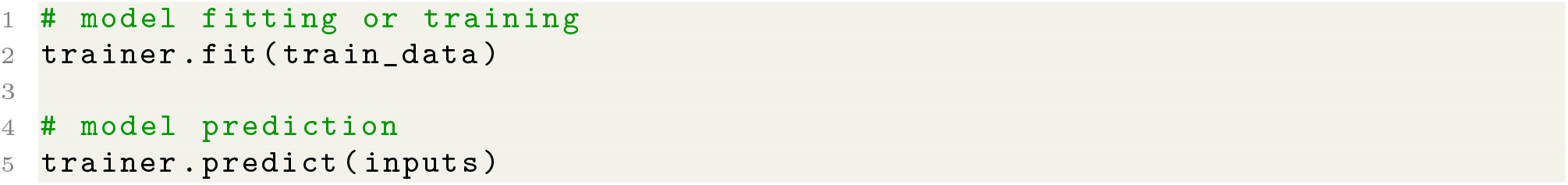

### 4.6 Analysis for brain dynamics models in BrainPy

The automatic analysis toolkit in BrainPy for low-dimensional dynamical systems relies on two numerical optimization methods, i.e., Brent’s method and BFGS algorithm. Given a one-dimensional system *dx/dt* = *f* (*x,t*), we want to find out its fixed points *f* (*x*\*,*T*) = 0 within the bound of x ∈ [*a, b*]. First, we screen out all intervals [*x*_1_,*x*_2_] that satisfy *f* (*x*_1_,*t*) * *f* (*x*_2_, *t*) ≤ 0. According to the intermediate value theorem, there must be a solution between *x*_1_ and *x*_2_ when *f* (*x*_1_) * *f* (*x*_2_) ≤ 0. Then we perform the Brent optimization in [*x*_1_,*x*_2_] to find out the root that satisfy *f* (*x*\*) = 0. Further, after obtaining the values of roots, BrainPy uses automatic differentiation to evaluate the stability of each root solution. In practice, users perform the phase plane analysis of a 1D system with

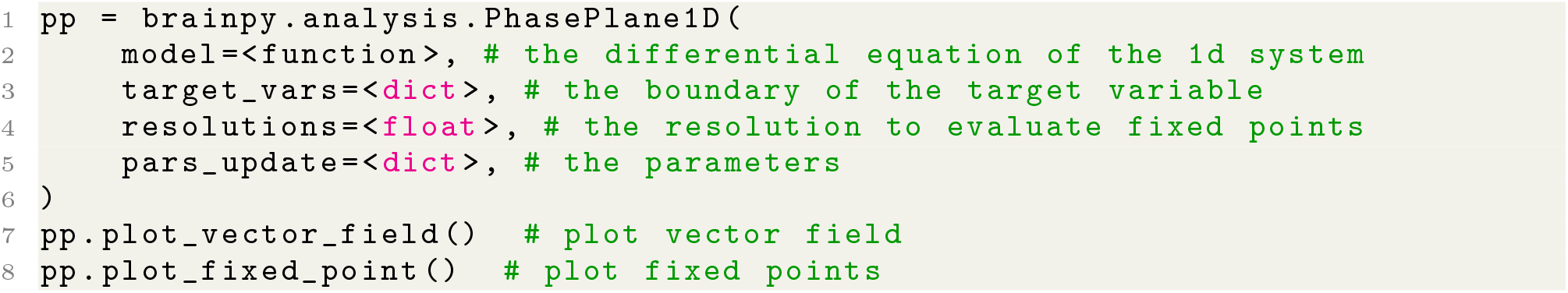

The bifurcation analysis of a 1D system is similar to its phase plane analysis, except specifying the target bifurcation parameters:

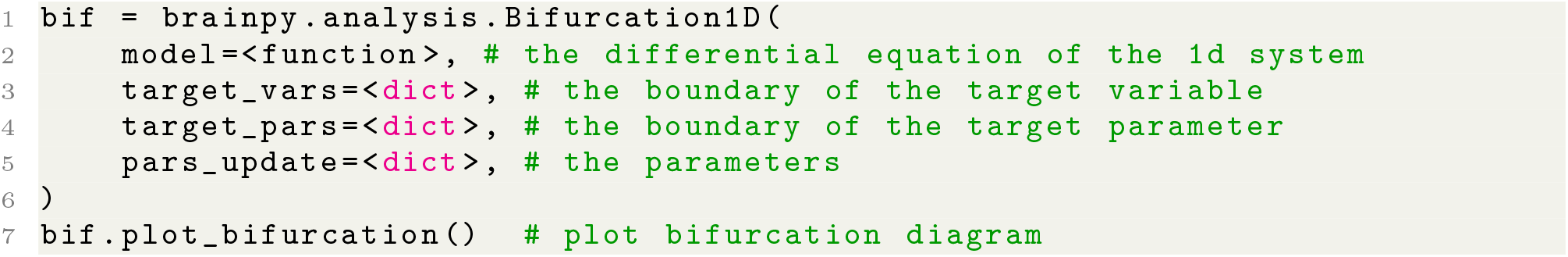

For a two-dimensional dynamical system 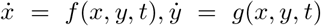, BrainPy performs its automatic analysis by defining an auxiliary function 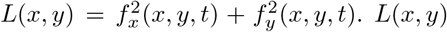 is always greater than zero, unless on its fixed points. Therefore, we utilize BFGS optimization to get all local minima. Then, we filter out the minima (*x*\*,*y*\*) whose losses *L*(*x*\*,*y*\*) are smaller than `tolerance`, typically, 1e^-8^, and we choose them as fixed points. Same as the one-dimensional system, we evaluate the stability of fixed points using automatic differentiation. In practice, users can perform phase plane and bifurcation analyses of a two-dimensional system with:

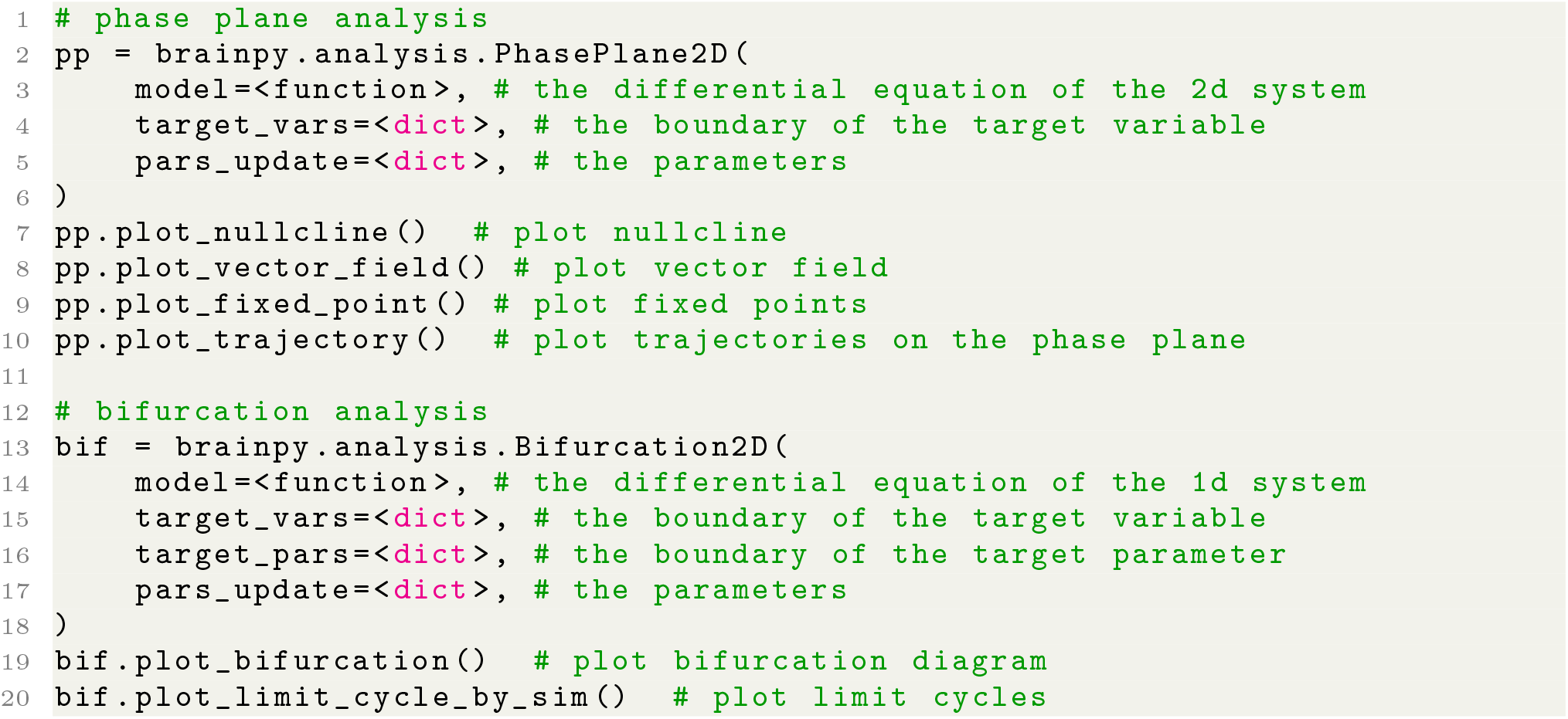

Similar to the analysis method of a two-dimensional system, we perform automatic analysis of a high-dimensional system with an auxiliary scalar function *L*(*X*). For the continuous system 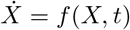, we use *L*(*X*) = |*f*(*X,t*)|^2^; for the discrete system *X*_*n*-1_ = *f* (*X*_*n*-1_,*t*), we use *L*(*X*) = |*X* – *f*(*X, t*)|^2^. If *X*\* is a fixed point, *L*(*X*\*) ≈ 0. Then, by incorporating the optimization method, including gradient descent method and nonlinear optimization algorithm like BFGS, we minimize the scalar function *L*(*X*) to search out the candidate fixed points. We finally select the fixed points by filtering out the point *X*\* that satisfies *L*(*X*\*) ≤ tolerance (typically, tolerance is 1e^-8^). In the later linearization analysis, we compute the Jacobian matrix at the fixed points, and evaluate the stability of a fixed point by the eigenvalues of its Jacobian matrix. In practice, after creating an instance of `DynamicalSystem`, users can perform its analysis by

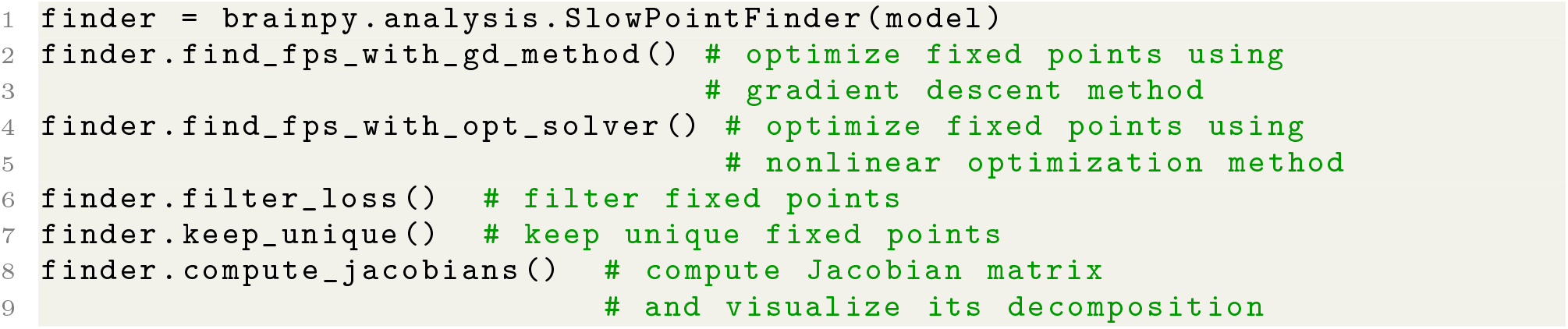

### 4.7 Extension of low-level operators in BrainPy

By bridging Numba [26], JAX [24] and XLA [25], BrainPy enables the customization of primitive operators through the native Python syntax. Exposing a custom operator to JAX requires registering an XLA “custom call”, and providing its C callback for Python. Based on the following two properties of Numba, we are aware of the possibility of using Numba as a convenient method for writing low-level kernels that support JAX’s JIT compilation. First, unlike JAX, which only supports JIT compilation of high-level functions, Numba is a JIT compiler that allows users to write a function with low-level fine-grained operations, for instance, looping over an array, or conditional branching over array values. This implies that `numba.jit()` can be used as a mean to write low-level kernel functions. The second property of Numba is that it provides a mechanism to create a compiled function that is callable from the foreign C code, such as XLA. Specifically, `numba.cfunc()` can be used to create a C callback for Numba’s JIT function to interface with XLA. Therefore, by integrating Numba with JAX and XLA, BrainPy provides an interface where users write primitive operators directly with Python syntax. Note that Numba supports various native Python features and a large subset of NumPy functions. Therefore, there is large flexibility in coding low-level primitives with Numba.

Below, we illustrate how to write a primitive operator in BrainPy. Particularly, to customize a primitive operator we need to provide two functions. The first is an abstract evaluation function that tells JAX what shapes and types of outputs are produced according to the input shapes and types:

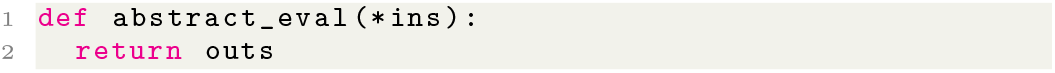

in which `ins` specifies the information of input shapes and types, `outs` denotes the array information of shapes and types of outputs. The other function is the concrete computation function, in which the output data is calculated according to the input data:

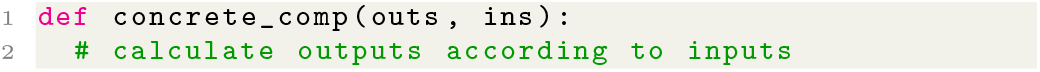

where `outs` and `ins` are output and input data owned by XLA, respectively. Note this function should not return anything. All computation must be made in-place. Finally, by using

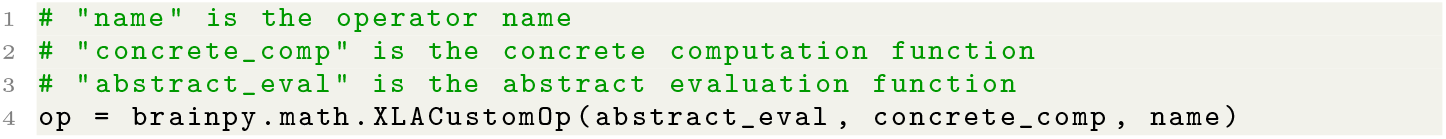

we register a primitive operator `op`. This operator `op` can be used in anywhere the user want.

## 5 Data availability

The preproceesed datasets are conveniently accessible in Python via the `brainpy.datasets` module. The data required to generate the figures and analyses in the paper are available at https://github.com/PKU-NIP-Lab/brainpy-paper-reproducibility/.

## 6 Code availability

BrainPy is distributed via the pypi package index (https://pypi.org/project/brainpy/) and is publicly released on GitHub (https://github.com/PKU-NIP-Lab/BrainPy/) under the license of GNU General Public License v3.0. Its documentation is hosted on the free documentation hosting platform *Read the Docs* (https://brainpy.readthedocs.io/). Rich examples and illustrations of BrainPy are publicly available at the website of https://brainpy-examples.readthedocs.io/. The source codes of these examples are available at https://github.com/PKU-NIP-Lab/BrainPyExamples/. All the codes to reproduce the results in the paper can be found at the following GitHub repository: https://github.com/PKU-NIP-Lab/brainpy-paper-reproducibility/.

## 7 Acknowledgements

This work was supported by Science and Technology Innovation 2030-Brain Science and Brain-inspired Intelligence Project (No. 2021ZD0200204) and Beijing Academy of Artificial Intelligence. We would like to acknowledge Xiaolong Zou, Xiaohan Lin, and all other members of the Wu laboratory for helpful discussions. CMW would like to acknowledge his family for their meticulous supports.

