## Supplemental Information for "*BrainPy*: a flexible, integrative, efficient, and extensible framework towards general-purpose brain dynamics programming"

**1 Background** **3**

**4 Supporting Tables** **24**

**5 Supporting Figures** **26**

**6 Supporting Codes** **30**

### 1 Background

#### 1.1 Review of the existing programming paradigm

In general, the existing tools for brain dynamics programming can be roughly divided into two categories: low-level programming and descriptive language.

The representatives of the first category include NEURON [1], NEST [2], CARLsim [3, 4], NeuronGPU [5], Arbor [6], and others. These simulators offer a library of standard models written in C or CUDA to guarantee efficient running, along with the convenient Python interface exposed to users for ease of use. Users can create neural networks in Python by grouping neuron and synapse models provided in the library. However, once a new model is needed, users must learn to program using the low-level language. This increases the learning cost dramatically and restricts the flexibility of defining new models [7].

The second category tools include Brian [8], Brian2 [9, 10], ANNarchy [11], GeNN [12], BMTK [13], NetPyNE [14], and NeuroML [15], which employ a code generation approach based on descriptive languages. Descriptive simulators allow users to create new models based on convenient descriptions, such as text [8–11], JSON [13, 14], or XML [15] files, and then translate the descriptions into low-level codes to speed up model running. In such a way, descriptive simulators enable model customization based on high-level descriptive languages and ensure efficient running by generating low-level codes.

Currently, descriptive language has become a standard approach for brain simulation [16]. Simulators employing the first approach also starts to provide their descriptive language interface for code generation. For example, NEST simulator provided its domain-specific language NESTML [17] to describe stereotypical neuron and synapse models. NEURON simulator recently released its modern descriptive interface NetPyNE [14], which employs the standard JSON format to allow users to describe neural circuit models by composing the existing available NEURON building block models. However, the customization of a new model for channels, neurons, or synapses still needs users to code based on its low-level programming interface.

Descriptive languages have achieved great success on brain simulations [16]. A major benefit of descriptive simulators is that they greatly simplify the modeling process by eliminating the need to manually write low-level efficient codes for a new dynamical model. However, they have intrinsic limitations on transparency, extensibility, and flexibility. One prominent feature of these descriptive languages is that they completely separate the model definition from the simulation, and therefore are not directly executable [16]. This kind of programming paradigm will cause great restrictions on usability and flexibility, because it disables the model debugging, error correction, and direct logic controlling. Moreover, descriptive languages are usually designed for specific kinds of models or one particular modeling approach. They are written in more than two programming languages: one is based on the low-level language (e.g., C++, CUDA) to implement its core functionality, the other is based on the high-level language (e.g., Python, Matlab) for ease of use. Once they are not tailored to users' needs, extensions to accommodate new components must be made in both high- and low-level languages, which is hard or nearly impossible for normal users. What's more, descriptive languages greatly reduce the the expressive power of a general-purpose programming language, and are hard to describe all aspects of a simulation experiment, including clipping variables out of bounds, input-output relations, model debugging, code optimization, dynamics analysis, and others.

In summary, great challenges on transparency, flexibility, efficiency, and extensibility are still present in the existing programming paradigm for brain simulation. We can draw the conclusion that current software solutions cannot lead us to a general-purpose programming framework that allows us to freely define brain dynamics models in various application domains.

#### 1.2 JIT compilation and JIT compilers: JAX, XLA and Numba

A notorious challenge in scientific computing is the trade-off between usability and efficiency. The former seeks to the fast-prototyping of thoughts and ideas, whereas the latter pursuits the efficient code execution. For a long time in the past, it was difficult to strike a balance between the two. For example, statically-typed compiled programming languages such as C or C++ are incredibly efficient in code execution, but their productivity is relatively low due to its complex and heavy syntax. In contrast, dynamically-typed interpreted programming languages like Python and R are easy to learn and use, but they have slow running speeds. Nowadays, with the increasing complexity of models, the demand for both usability and efficiency has increased dramatically. Fortunately, recent advancements in just-in-time (JIT) compilation technology [18, 19] have provided viable answers to this two language problem. In particular, a new generation of computational engines based on JIT compilation [20] has begun to have an impact in a variety of scientific computing disciplines.

Intuitively speaking, JIT compilation is the union of the statically-typed compilation and dynamically-typed interpretation. It takes advantage of both the convenience of dynamic high-level languages like Python and the efficiency of static low-level languages such as C++. At the beginning of a program running, a JIT compiler behaves like an interpreter. It runs your code step by step, and is able to output the intermediate results at the run-time for debugging. However, if some hot code snippets that executed frequently, e.g., certain functions or loop bodies, are detected or manually labeled, they will be submitted to the JIT compiler for compilation and storage. In this sense, it behaves like a statically-typed compiler. Once the compiled code snippets are entered again, the program will directly execute the compiler-generated low-level code without time-consuming interpreting again. Hot code snippets can be automatically detected by the JIT compiler, or be manually labeled by users.

JIT compilation has been a mature and well-accepted technology. It has been adopted in modern programming languages like JAVA, Julia, and Python. JAVA language provides JIT compilation in its JAVA virtual machine (JVM) to accelerate the execution of JAVA code [21]. Java source code is first compiled into the platform-independent Java bytecode (`.class` file). Then, JVM loads the bytecode, interprets it, and executes it. To increase the running speed, JVM detects code that are frequently called through the hotspot detection and submits their bytecode to the JIT compiler to compile them into machine code. For the code with lower frequency, executing it through the JVM interpretation can save the time of the JIT compilation; while for the hot code frequently called, JIT compilation can significantly improve the running speed after the code is compiled. However, compared with Python, JAVA has poor ecosystem supports on numerical computing. Its JIT is not focused on numerical computing, but for general domains.

Recently, Julia [22], another dynamic high-level programming language, is proposed for high-performance scientific computing. Julia features the intuitive, productive and general-purpose syntax which inspired from the success of Python, Matlab, and C++. Meanwhile, it achieves the attractive performance through the JIT compilation based on the LLVM compiler infrastructure [18]. In an amazing short time, Julia has offered wonderful routines for mathematical functions, machine learning algorithms, data processing tools, visualization utilities, and others. However, Julia is still young. Costs, like lack of familiarity, rough edges, correctness bugs, and continual language changes, are still imposed to normal users.

Python is an well-known and popular interactive dynamic programming language. It has a long history on numerical computing [23, 24]. Ecosystem on scientific computing, including array programming [24], scientific algorithms [25], machine learning [26], deep learning [27–29], image processing [30], data analysis and statistics [31], network analysis [32], visualization [33], and many others, has been well established in Python. Before Julia, JIT compilation was introduced into Python by PyPy since 2007. Later, other attempts, including Pyston, Pyjion, Psyco, JitPy, HOPE, et. al, are proposed. With a long history on JIT development, Python nowadays has provided mature platforms of JIT compilation focusing on numerical computing. These numerical JIT platforms include Numba [34], JAX [29], and XLA [35]. Each of them has its own characteristics.

JAX [29] is a flourishing machine learning library developed by Google. It aims to provide high-level numerical functions to help users fast prototype machine learning ideas. Moreover, these numerical functions can benefit from the powerful functional transformations, like automatic differentiation `grad`, JIT compilation `jit`, automatic vectorization `vmap`, and parallelization `pmap`. JAX makes a heavy use of XLA [35] (see the following text) for code optimization. Specifically, for ease of use, high-level numerical functions in JAX are NumPy-like. JAX provides many numerical functions in NumPy, including basic mathematical operators, linear algebra functions, and Fourier transform routines (see Table 1). However, some fundamental designs are significantly different from NumPy, for instance, the well-established syntax for in-place updating and random samplings. This is the reason why we provide another set of numerical functions consistent with NumPy (see main text). In addition to its NumPy-like API, JAX provides a wonderful set of composable functional transformations. Among them, automatic differentiation in JAX supports both forward and backward modes for arbitrary numerical functions. It can take derivatives of a function with a large subset of Python syntax, including loops, conditions, recursions, and closures. Moreover, JAX utilizes XLA to just-in-time compile your Python code on modern devices, like CPUs, GPUs and TPUs. It can significantly accelerate the execution speed of your code, and allows you to get maximal performance without having to leave Python. JAX also provides automatic vectorization or batching. It supports to transform loops to vector operations via a single functional call `vmap`. What's more, JAX delivers `pmap` to express single-instruction multiple-data (SIMD) programs. Applying `pmap` to a function will just-in-time compile and execute the code in parallel on XLA devices, like multiple GPUs or TPU cores. Similar with `vmap`, `pmap` transformation maps a function over array axes. But what's different is that the former vectorizes functions by compiling the mapped axis as primitive operations, whereas the later replicates the function and runs each replica on its own XLA device in parallel. Automatic differentiation and compilation in JAX can be composed arbitrarily to enable rapid experimentation of novel algorithms.

XLA [35] is a domain specific linear algebra compiler developed by Google which aims to improve the execution speed, memory usage, portability, and mobile footprint reduction of machine learning algorithms. XLA compiler provides the support for JIT compilation based on LLVM [18]. The front end program (e.g., JAX) which want to take advantage of JIT compilation of XLA should first define the computation graph as "High Level Optimizer IR" (HLO IR). Then, XLA takes this graph defined in HLO IR and compiles it into machine instructions for different backend architectures. Currently, XLA supports JIT compilation on backend devices of x86-64 CPUs, NVIDIA GPUs, and Google TPUs. XLA is designed for easy portability on new hardware. It provides an abstract interface that a new hardware device can implement to create a backend to run existing computation graphs. Instead of implementing every existing operator for new hardware, XLA provides a simple and scalable mechanism which can retarget different backend. This advantage may be valuable for neuromorphic computing [36], because new neuromorphic hardware can be interfaced as a new backend of XLA computation.

Numba [34] is a JIT compiler for numerical functions in Python. Similar with JAX, Numba supports JIT compilation of a subset of Python and NumPy code based on LLVM compiler. However, different from JAX which accelerates the computation flow composed of high-level operators, Numba pays more attention on the acceleration of loop-based functions. Numba achieves excellent optimizations on Python functions with a lot of loops. It allows users to write fine-grained code with native Python control flows and meanwhile obtain the running speed approaching C. This is a huge advantage compared to JAX, because JAX does not support the automatic vectorization of a for-loop function.

### 2 Materials and Methods

#### 2.1 Continuous integration and documentation generation

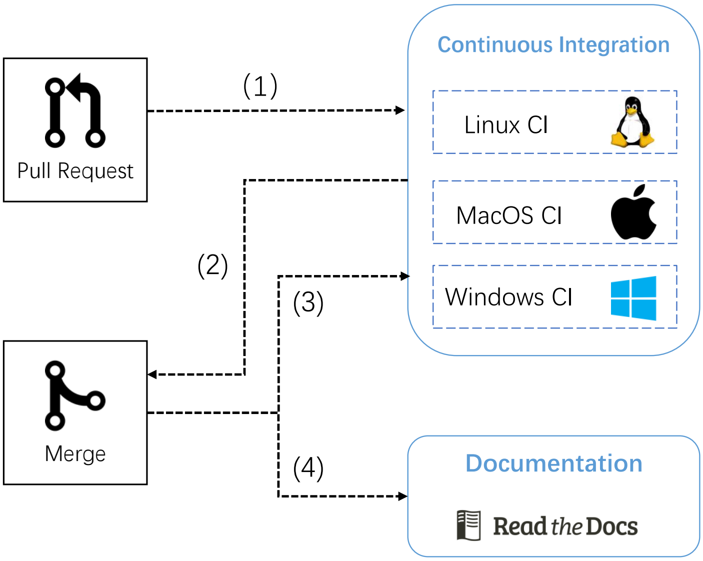

**SI Fig. 1 : The pipeline of automatic continuous integration and documentation building in BrainPy.**

To ensure any code changes do not introduce unintended bugs or side-effects, we standardized the development process and enabled the automatic continuous integration (CI) of BrainPy with GitHub Actions. Moreover, to update the tutorial and documentation with the latest code changing, we automated the documentation building of BrainPy with *Read the Docs*. SI Fig. 1 illustrates the whole workflow of BrainPy development. First, any code change should be proposed through GitHub Pull Request. Once a Pull Request is opened, CI pipelines are triggered to test BrainPy codes on Windows, Linux and MacOs platforms (SI Fig. 1 (1)). After all test suites are passed, the code reviewer should manually inspect the significance and correctness of the proposed code changes again. If all things are fine, the reviewer can merge the code into the master branch (SI Fig. 1 (2)). After merging, a new set of test cases are triggered automatically to test the latest BrainPy codebase doesn't have bugs (SI Fig. 1 (3)). Besides, the merging operation also triggers the automatic documentation generation through the documentation hosting platform *Read the Docs* (SI Fig. 1 (4)), in which the latest documentation, including the code annotation, user manual and tutorials, are automatically built with Sphinx and hosted online at https://brainpy.readthedocs.io/.

##### 2.2 Selection of brain simulators for comparison

To identify the state-of-art brain simulators for comparison, we follow the approach of Tikidji-Hamburyan et al. [7], and analyze the most popular simulation environments registered in *ModelDB* database: https://senselab.med.yale.edu/ModelDB/FindBySimulator.cshtml. In total, there are 116 simulation environments registered in *ModelDB*. We remove the records using general-purpose programming languages (e.g., Python, Matlab, C++, Julia, R), clean simulators with zero number of records, and merge duplicate simulator names in the list. Then, there are 83 software left. Among them, we quantify the popularity index of a simulator by counting its number of records publicly available in the *ModelDB* website. The five most popular simulators according to the popularity index rank are NEURON [1], Brian [8, 10], GENESIS [37], SNNAP [38], and NEST [2]. However, due to GENESIS [37] has no longer been maintained and SNNAP [38] has a poor documentation, we remove both of them out of the comparison. Finally, our comparison list for brain simulators include NEURON, NEST, and Brian2.

#### 3 Supplemental Results

##### 3.1 The programming interface for neuron models

For reduced neuron models, like integrate-and-fire model [39] and Izhkevich neuron model [40], we can directly define them as `DynamicalSystem` classes (see main text). As for Hodgkin–Huxley typed neuron models, which contain multiple ion channels, BrainPy provides a convenient interface for their definition.

In general, a Hodgkin–Huxley typed neuron model is characterized by the following equation:

$$C_m \frac{dV}{dt} = \sum_j g_j (E - V) + I_{ext}, \tag{1}$$

where $g_j = \bar{g}_j M^x N^y$ is the channel conductance, $E$ is the reversal potential, $M$ is the activation variable, and $N$ is the inactivation variable.

$M$ and $N$ have the dynamics of

$$\frac{dx}{dt} = \phi_x \frac{x_\infty(V) - x}{\tau_x(V)}, \tag{2}$$

where $x \in [M, N]$, $\phi_x$ is a temperature-dependent factor, $x_\infty$ is the steady state, and $\tau_x$ is the time constant. Equivalently, the above equation can be rewritten as:

$$\frac{dx}{dt} = \phi_x \left(\alpha_x(1-x) - \beta_x x\right) \tag{3}$$

where $\alpha_x$ and $\beta_x$ are rate constants.

For this kind of Hodgkin–Huxley typed neuron models, BrainPy provides `brainpy.dyn.CondNeuGroup` to model its dynamics. Specifically, the ion channel models added into the `brainpy.dyn.CondNeuGroup` instance can be automatically evaluated and updated.

Let's take a model of high-threshold thalamocortical (HTC) cell [41] as the example. This HTC model contains ten ion channel currents: a leakage channel current ($I_L$), a potassium leak channel current ($I_{KL}$) to mimic the neuromodulation effect, a spike generating fast sodium current ($I_{Na}$), a delayed rectifier potassium current ($I_{DR}$), a hyperpolarization-activated cation current ($I_H$), a high-threshold L-type $Ca^{2+}$ current ($I_{Ca/L}$), a $Ca^{2+}$-dependent potassium current ($I_{AHP}$), a $Ca^{2+}$-activated nonselective cation current ($I_{CAN}$), a regular low-threshold T-type $Ca^{2+}$ current ($I_{Ca/T}$), and a high-threshold T-type $Ca^{2+}$ current ($I_{Ca/HT}$). All these ionic currents can be easily gathered with the BrainPy syntax to form a conductance-based neuron model.

```
import brainpy as bp

class HTC(bp.dyn.CondNeuGroup):
  def __init__(self, size, gKL=0.01, gL=0.01):
    IL = bp.channels.IL(size, g_max=gL, E=-70)
    IKL = bp.channels.IKL(size, g_max=gKL)
    INa = bp.channels.INa_Ba2002(size, V_sh=-30)
    IDR = bp.channels.IKDR_Ba2002(size, V_sh=-30., phi=0.25)
    Ih = bp.channels.Ih_HM1992(size, g_max=0.01, E=-43)

    # calcium dependent channels
    ICaL = bp.channels.ICaL_IS2008(size, g_max=0.5)
    IAHP = bp.channels.IAHP_De1994(size, g_max=0.3, E=-90.)
    ICaN = bp.channels.ICaN_IS2008(size, g_max=0.5)
    ICaT = bp.channels.ICaT_HM1992(size, g_max=2.1)
    ICaHT = bp.channels.ICaHT_HM1992(size, g_max=3.0)
    Ca = bp.channels.CalciumDetailed(size, C_rest=5e-5, tau=10., d=0.5,
                                     IAHP=IAHP, ICaN=ICaN, ICaT=ICaT,
                                     ICaL=ICaL, ICaHT=ICaHT)

    super(HTC, self).__init__(size, A=2.9e-4, V_th=20.,
                              IL=IL, IKL=IKL, INa=INa,
                              IDR=IDR, Ih=Ih, Ca=Ca)
```

#### 3.2 The programming interface for synapse models

In BrainPy, we provide a general class `brainpy.dyn.SynConn` to help users to model any synaptic model. Moreover, we provide `brainpy.dyn.TwoEndConn` to model the synaptic projection between pre- and post-synaptic neuron groups.

`brainpy.dyn.TwoEndConn` can be used to model synapse models with the form of:

$$
\begin{aligned}
\frac{\mathrm{d}g(t)}{\mathrm{d}t} &= f_{\mathrm{dyn}}(g(t), t) && \rightarrow \text{synaptic dynamics} && (4)\\
g_{\max}(t) &= f_{\mathrm{LTP}}(g_{\max}(t), t) && \rightarrow \text{synaptic long-term plasticity} && (5)\\
g(t) &= f_{\mathrm{STP}}(g(t), t) && \rightarrow \text{synaptic short-term plasticity} && (6)\\
I_{\mathrm{post}}(t) &= f_{\mathrm{out}}(g_{\max}(t) \cdot g(t), t) && \rightarrow \text{synaptic outputs} && (7)\\
& && && (8)
\end{aligned}
$$

in which

- $g \in [0, 1]$ is the synaptic conductance which represents the fraction of the open channels.
- $g_{\max}$ is the synaptic weight,
- $I_{\mathrm{post}}$ is the synaptic current onto the post-synaptic neuron group,
- $f_{\mathrm{dyn}}$ is the function to compute synaptic dynamics,
- $f_{\mathrm{LTP}}$ is the function for computing synaptic long-term plasticity,
- $f_{\mathrm{STP}}$ is the function for computing synaptic short-term plasticity,
- $f_{\mathrm{out}}$ is the way to output synaptic currents onto post-synaptic neurons.

For instance, an exponential synapse model has the dynamics $f_{\mathrm{dyn}}$ of

$$\frac{\mathrm{d}g}{\mathrm{d}t} = -\frac{g}{\tau_{\mathrm{decay}}} + \sum_k \delta(t - t^k), \tag{9}$$

a dual exponential synapse model has the dynamics $f_{\mathrm{dyn}}$ with the form of

$$\frac{\mathrm{d}g}{\mathrm{d}t} = -\frac{g}{\tau_{\mathrm{decay}}} + h, \tag{10}$$

$$\frac{\mathrm{d}h}{\mathrm{d}t} = -\frac{h}{\tau_{\mathrm{rise}}} + \sum_k \delta\left(t - t^{(f)}\right). \tag{11}$$

Both two models can output synaptic currents with the current-based form:

$$I = g_{\max} \cdot g(t), \tag{12}$$

or the conductance-based form:

$$I = g_{\max} \cdot g(t) \cdot (V_{\mathrm{post}}(t) - E). \tag{13}$$

Moreover, the synapse conductance $g$ can be filtered by short-term plasticity variables $u, x$ [42]. For current-based synapses,

$$I = g_{\max} \cdot g(t) u x, \tag{14}$$

for conductance-based synapses,

$$I = g_{\max} \cdot g(t)ux \cdot (V_{\text{post}}(t) - E). \tag{15}$$

What's more, the synapse weight $g_{\max}$ can be adaptively changed by long-term plasticity rules, like Spike Timing Dependent Plasticity (STDP) [43].

All these combinations can be easily achieved with `brainpy.dyn.TwoEndConn` to implement various synapse models. For example, a current-based synapse model can be defined with:

```
import brainpy as bp

# with exponential decay dynamics
syn = bp.synapses.Exponential(..., output=bp.synouts.CUBA())

# with alpha dynamics
syn = bp.synapses.Alpha(..., output=bp.synouts.CUBA())

# with dual exponential decay dynamics
syn = bp.synapses.DualExponential(..., output=bp.synouts.CUBA())
```

A conductance-based synapse model can be defined with:

```
# with exponential decay dynamics, reversal potential is 0. mV
syn = bp.synapses.Exponential(..., output=bp.synouts.COBA(E=0.))

# with alpha dynamics
syn = bp.synapses.Alpha(..., output=bp.synouts.COBA())

# with dual exponential decay dynamics
syn = bp.synapses.DualExponential(..., output=bp.synouts.COBA())
```

Adding a short-term plasticity effect onto a synapse becomes very easy. For instance,

```
# exponential synapse model with STD
syn = bp.synapses.Exponential(..., output=bp.synouts.CUBA(),
                                   stp=bp.synplast.STD())

# AMPA synapse model with STP
syn = bp.synapses.AMPA(..., output=bp.synouts.COBA(),
                            stp=bp.synplast.STP())
```

Simulating the long-term plasticity effect in a synapse model is also a piece of cake. For instance,

```
# exponential synapse model with STDP
syn = bp.synapses.Exponential(..., output=bp.synouts.CUBA(),
                                   ltp=bp.synplast.STDP())

# AMPA synapse model with STDP
syn = bp.synapses.AMPA(..., output=bp.synouts.COBA(),
                            ltp=bp.synplast.STDP())
```

#### 3.3 The programming interface for network models

Typically, a network in BrainPy is decomposed into neuron groups and their synaptic projections. Taking an E/I balanced network [44] as the example, its structure is given as the following (see Fig. 2):

- a group of excitatory neurons ("E"),

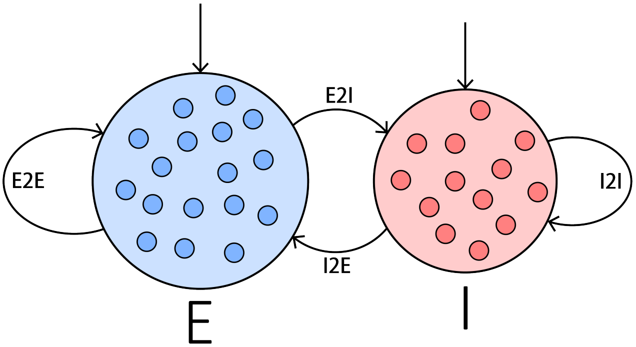

**SI Fig. 2** : The structure of an E/I balanced network model. "E" denotes the excitatory population, "I" the inhibitory population, "E2E" the self-connection within the E population, "I2I" the self-connection within the I population, "E2I" the synaptic projection from E population to I population, and "I2E" the synaptic projection from I population to E population.

- a group of inhibitory neurons ("I"),
- synaptic connections within the excitatory and inhibitory neuron groups, respectively ("E2E" and "I2I"), and
- the inter-connections between these two groups ("E2I" and "I2E").

Therefore, we can program an E/I balanced network model by inheriting from `brainpy.dyn.Network` (a subclass of `brainpy.dyn.DynamicalSystem` that is specifically used to create network models):

```
import brainpy as bp

class EINet(bp.dyn.Network):
  def __init__(self):
     self.E = YourNeuronModel(...)
     self.I = YourNeuronModel(...)
     self.E2E = YourSynapseModel(self.E, self.E, ...)
     self.E2I = YourSynapseModel(self.E, self.I, ...)
     self.I2E = YourSynapseModel(self.I, self.E, ...)
     self.I2I = YourSynapseModel(self.I, self.I, ...)
```

Note here we no longer need to overwrite `update()` function because `bp.dyn.Network` can automatically evaluate it.

As for a more complicated decision-making network model [45], which contains ten neuron groups and mutual projections between each other, defining it is just creating the corresponding sub-populations and their mutual connections (see SI Listing 1).

```
import brainpy as bp

class DecisionMakingNet(bp.dyn.Network):
  def __init__(self, mu0=40., coherence=25.6, f=0.15):
    super(DecisionMaking, self).__init__()

    num_A = num_B = int(f * 1600)
    num_N = 1600 - num_A - num_B
    num_inh = 400
```

```
# E neurons/pyramid neurons
self.A = bp.neurons.LIF(num_A, ...)
self.B = bp.neurons.LIF(num_B, ...)
self.N = bp.neurons.LIF(num_N, ...)

# I neurons/interneurons
self.I = bp.neurons.LIF(num_inh, ...)

# poisson stimulus
self.IA = PoissonStim(num_A, freq_mean=mu0 + mu0 / 100. * coherence)
self.IB = PoissonStim(num_B, freq_mean=mu0 - mu0 / 100. * coherence)

# noise neurons
self.noise_A = bp.neurons.PoissonGroup(num_A, freqs=2400.)
self.noise_B = bp.neurons.PoissonGroup(num_B, freqs=2400.)
self.noise_N = bp.neurons.PoissonGroup(num_N, freqs=2400.)
self.noise_I = bp.neurons.PoissonGroup(num_inh, freqs=2400.)

# define external inputs
self.IA2A = bp.synapses.Exponential(self.IA, self.A, ...)
self.IB2B = bp.synapses.Exponential(self.IB, self.B, ...)

# define E2E conn
self.A2A_AMPA = bp.synapses.Exponential(self.A, self.A, ...)
self.A2A_NMDA = bp.synapses.NMDA(self.A, self.A, ...)

self.A2B_AMPA = bp.synapses.Exponential(self.A, self.B, ...)
self.A2B_NMDA = bp.synapses.NMDA(self.A, self.B, ...)

self.A2N_AMPA = bp.synapses.Exponential(self.A, self.N, ...)
self.A2N_NMDA = bp.synapses.NMDA(self.A, self.N, ...)

self.B2A_AMPA = bp.synapses.Exponential(self.B, self.A, ...)
self.B2A_NMDA = bp.synapses.NMDA(self.B, self.A, ...)

self.B2B_AMPA = bp.synapses.Exponential(self.B, self.B, ...)
self.B2B_NMDA = bp.synapses.NMDA(self.B, self.B, ...)

self.B2N_AMPA = bp.synapses.Exponential(self.B, self.N, ...)
self.B2N_NMDA = bp.synapses.NMDA(self.B, self.N, ...)

self.N2A_AMPA = bp.synapses.Exponential(self.N, self.A, ...)
self.N2A_NMDA = bp.synapses.NMDA(self.N, self.A, ...)

self.N2B_AMPA = bp.synapses.Exponential(self.N, self.B, ...)
self.N2B_NMDA = bp.synapses.NMDA(self.N, self.B, ...)

self.N2N_AMPA = bp.synapses.Exponential(self.N, self.N, ...)
self.N2N_NMDA = bp.synapses.NMDA(self.N, self.N, ...)

# define E2I conn
self.A2I_AMPA = bp.synapses.Exponential(self.A, self.I, ...)
self.A2I_NMDA = bp.synapses.NMDA(self.A, self.I, ...)

self.B2I_AMPA = bp.synapses.Exponential(self.B, self.I, ...)
self.B2I_NMDA = bp.synapses.NMDA(self.B, self.I, ...)

self.N2I_AMPA = bp.synapses.Exponential(self.N, self.I, ...)
self.N2I_NMDA = bp.synapses.NMDA(self.N, self.I, ...)

# define I2E conn
self.I2A = bp.synapses.Exponential(self.I, self.A, ...)
self.I2B = bp.synapses.Exponential(self.I, self.B, ...)
self.I2N = bp.synapses.Exponential(self.I, self.N, ...)

# define I2I conn
self.I2I = bp.synapses.Exponential(self.I, self.I, ...)
```

```
    # define external projections
    self.noise2A = bp.synapses.Exponential(self.noise_A, self.A, ...)
    self.noise2B = bp.synapses.Exponential(self.noise_B, self.B, ...)
    self.noise2N = bp.synapses.Exponential(self.noise_N, self.N, ...)
    self.noise2I = bp.synapses.Exponential(self.noise_I, self.I, ...)
```

**SI Listing 1** : Building a decision-making network model [45] with `brainpy.dyn.Network` . This code snippet is just for illustration. For the complete source code to run this network, please see https://brainpy-examples.readthedocs.io/en/latest/decision_making/Wang_2002_decision_making_spiking.html.

#### 3.4 Ecosystem of brain models

BrainPy provides the modular and composable programming interface to define brain dynamics models. With the guidance of this programming paradigm, BrainPy provides rich brain models as building blocks to help fast-prototype high-level models. These building blocks include neuron models, synapse models, population rate models, ion channel models, reservoir computing models, artificial neural network models, and others.

Providing rich built-in brain dynamics models has several advantages:

1. Simplify the process of model building. Users without programming expertise can easily build complex brain dynamics models with these building blocks.
2. Examplify how to customize models. These models can be used as examples to illustrate how to customize models in BrainPy.
3. Standardize model implementations. These built-in models can also be used as standard models to reproduce simulation results, thus helping communication between investigators.

In the future, we will continue to update the models commonly used in computational neuroscience research.

##### 3.4.1 Neuron models

BrainPy provides multiple kinds of neuron models, including biological and reduced neurons models which are modeled with ODEs or SDEs, and fractional-order neuron models that are using FDEs. This kind of models include, but are not limited to:

- Biological neuron models
  - Hodgkin–Huxley model [46]
  - Morris-Lecar model [47]
  - Pinsky-Rinsel model [48]
  - Wang-Buzsaki model [49]
- Reduced neuron models
  - Leaky integrate-and-fire neuron model [39]
  - Exponential integrate-and-fire neuron model [50]

  - Adaptive exponential integrate-and-fire neuron model [50]
  - Quadratic Integrate-and-Fire neuron model [51]
  - Adaptive quadratic integrate-and-fire neuron model [52]
  - Generalized Integrate-and-Fire model [53]
  - Leaky Integrate-and-Fire model with SFA [54]
  - Izhikevich neuron model [40]
  - Hindmarsh-Rose neuron model [55]
  - FitzHugh-Nagumo neuron model [56]
  - Reduced TRN neuron model [57]

- Fractional-order neuron models
  - Fractional-order FitzHugh-Rinzel model [58]
  - Fractional-order Izhikevich model [59]

##### 3.4.2 Synapse models

BrainPy provides chemical synapse models, electrical synapse models, and plasticity models. This kind of models include, but are not limited to:

- Chemical synapse models
  - Delta synapse model
  - Exponential decay synapse model [60]
  - Dual exponential synapse model [60]
  - Alpha synapse model [60]
  - NMDA synapse model [61]
  - AMPA synapse model [62]
  - $\mathrm{GABA_A}$ synapse model [63]
  - $\mathrm{GABA_B}$ synapse model [64]
- Electrical synapse models
  - Diffusive coupling model
  - Additive coupling model
  - Gap junction model [65]
- Plasticity synapse models
  - Short-term plasticity model [42]
  - Long-term plasticity model [66]

##### 3.4.3 Population rate models

BrainPy also provides a series of population rate models which describe the collective behavior of a poulation of neurons. This kind of models include, but are not limited to:

- FitzHugh-Nagumo model [67]
- Feedback FitzHugh-Nagumo model [68]
- Quadratic integrate-and-fire population model [69]

- Stuart-Landau population model [70, 71]
- Wilson-Cowan population model [72]
- Threshold linear rate model [73]
- Theta neuron model [74]
- Jansen-Rit model [75]
- Van der Pol oscillator [76]

##### 3.4.4 Ion channel models

Neurons are typical made up of ion channels. BrainPy provides `brainpy.dyn.CondNeuGroup` for constructing conductance-based neuron models by composing ion channels. Our built-in ion channel models include voltage-dependent Sodium channel models, voltage-dependent Potassium channel models, voltage-dependent Calcium channel models, Calcium-dependent Potassium channel models, hyperpolarization-activated Cation channel models, and leakage channel models. This kind of models include, but are not limited to:

- Voltage-dependent Sodium channel models
  - `brainpy.channels.INa_Ba2002` [77]
  - `brainpy.channels.INa_TM1991` [78]
  - `brainpy.channels.INa_HH1952` [46]
- Voltage-dependent Potassium channel models
  - `brainpy.channels.IKDR_Ba2002` [77]
  - `brainpy.channels.IK_TM1991` [78]
  - `brainpy.channels.IK_HH1952` [46]
  - `brainpy.channels.IKA1_HM1992` [79]
  - `brainpy.channels.IKA2_HM1992` [79]
  - `brainpy.channels.IKK2A_HM1992` [79]
  - `brainpy.channels.IKK2B_HM1992` [79]
  - `brainpy.channels.IKNI_Ya1989` [80]
- Voltage-dependent Calcium channel models
  - `brainpy.channels.CalciumDetailed` [81, 82]
  - `brainpy.channels.ICaN_IS2008` [83]
  - `brainpy.channels.ICaT_HM1992` [79]
  - `brainpy.channels.ICaT_HP1992` [84]
  - `brainpy.channels.ICaHT_HM1992` [79]
  - `brainpy.channels.ICaL_IS2008` [85]
- Calcium-dependent Potassium channel models
  - `brainpy.channels.IAHP_De1994` [83]
- Hyperpolarization-activated Cation channel models
  - `brainpy.channels.Ih_HM1992` [79]

  - `brainpy.channels.Ih_De1996` [86]

- Leakage channel models
  - The leakage channel current
  - The potassium leak channel current

##### 3.4.5 Reservoir computing models

Reservoir models have been useful tools in understanding how the brain works [87–89]. BrainPy provides several models for constructing classical reservoir computing models, like echo state machines [90], and next-generation reservoir computing models [91]. This kind of models include, but are not limited to:

- Nonlinear vector autoregression machine [91]
- Reservoir layer [90]

##### 3.4.6 Artificial neural networks

BrainPy also provides neural network models come from deep learning community. This kind of models include, but are not limited to:

- Conv1D [92]
- Conv2D [92]
- Conv3D [92]
- BatchNormal1D [93]
- BatchNormal2D [93]
- BatchNormal3D [93]
- Dropout [94]
- AveragePooling
- MaxPooling
- MinPooling
- Dense
- VanillaRNN
- GRU [95]
- LSTM [96]

#### 3.5 Parameter exploration and selection with parallel simulation

Parameter exploration and selection is an unavoidable process in brain dynamics modeling. BrainPy supports multiple kinds of approaches for parameter exploration.

Technically, parameter exploration requires parallelization, because it involves simulating multiple instances of the model with different parameter settings. BrainPy supports parallelization of multi-threading and multiprocessing on a single machine, and parallelization across multiple devices.

##### 3.5.1 Thread-based parallelization

Multi-threaded running of BrainPy models can be easily achieved.

The first approach is directly using the Python's multi-threading support. For example, SI Listing 2 demonstrates that by utilizing the `threading` backend of `joblib` library, we can easily achieve parallel execution based on multiple Python threads. However, the multi-threading parallelization based on this approach will get stuck in the well-known issue of Global Interpreter Lock (GIL) of Python.

```
from joblib import Parallel, delayed, parallel_backend

def run_model(par):
  model = YourModel(par)
  runner = bp.dyn.DSRunner(model)
  runner.run(<int>)
  return runner.mon

# define all parameter values need to explore
all_params = [...]

# create a multi-threading environment for batch simulation
with parallel_backend(backend="threading"):
  r = Parallel()([delayed(run_model)(p) for p in all_params])
```

**SI Listing 2** : The multi-threading parallelization of BrainPy models with `joblib`. `par` is a specific setting of the model parameter. `YourModel` defines your brain dynamics model. A full example of multi-threading parallelization based on `joblib` please refers to SI Listing 10.

The second approach of realizing multi-threading parallelization is the vectorization map of JAX `jax.vmap`. `jax.vmap` vectorizes functions by compiling the mapped axis as primitive operations. It can avoid the recompilation of models in the same batch, and automatically parallelize the model running on the given machine. Different from the first approach, the multi-threading parallelization of `jax.vmap` is implemented outside of the Python interpreter, so that the GIL problem no longer exists. SI Listing 3 demonstrates how simple of this parallelization approach is.

```
from jax import vmap

def run_model(par):
  model = YourModel(par)
  runner = bp.dyn.DSRunner(model)
  runner.run(<int>)
  return runner.mon

# define all parameter values need to explore
all_params = [...]

# batch simulation through jax.vmap
r = vmap(run_model)(*all_params)
```

**SI Listing 3** : The multi-threading parallelization of BrainPy models with `jax.vmap`. `par` is a specific setting of the model parameter. `YourModel` is your defined model. A full example of multi-threading parallelization based on `jax.vmap` please refers to SI Listing 11.

##### 3.5.2 Processor-based parallelization

Another way to avoid the Python GIL problem is the multi-processing parallelization. Based on `multiprocessing` library or `joblib` package, we can easily run multiple models concurrently on separate Python worker processes.

```
from joblib import Parallel, delayed, parallel_backend

def run_model(par):
  model = YourModel(par)
  runner = bp.dyn.DSRunner(model)
  runner.run(<int>)
  return runner.mon

# define all parameter values need to explore
all_params = [...]

# create a multi-processing environment for parallel simulation
with parallel_backend(backend="loky"):
  r = Parallel()([delayed(run_model)(p) for p in all_params])
```

**SI Listing 4** : The multi-processing parallelization of BrainPy models with `joblib`. What different from the code of SI Listing 2 is the `backend` is set to "loky". A full example of multi-processing parallelization based on `joblib` please refers to SI Listing 12.

##### 3.5.3 Multi-device parallelization

BrainPy models also intrinsically support the parallelization running on multiple devices (e.g., multiple GPU devices or TPU cores) or HPC systems (e.g., supercomputers). Different from the above thread-based and processor-based parallelization methods, in which the same model runs in parallel in different parts of memory on the same device, device-based parallelization runs the same model in parallel on multiple devices.

One way to express the multi-device parallelization of BrainPy models is using `jax.pmap` instruction. JAX delivers `jax.pmap` to express SIMD programs. It provides an interface to run the same model on multiple devices with different parameter values. It usage is analogy to `jax.vmap`. SI Listing 5 presents an example to run BrainPy models on multiple devices.

```
from jax import pmap

def run_model(par):
  model = YourModel(par)
  runner = bp.dyn.DSRunner(model)
  runner.run(<int>)
  return runner.mon

# define all parameter values need to explore
all_params = [...]

# parallel simulation through jax.pmap
r = pmap(run_model)(*all_params)
```

**SI Listing 5** : The multi-device parallelization of BrainPy models with `jax.pmap`. A full example of multi-device parallelization based on `jax.pmap` please refers to SI Listing 13.

BrainPy also works well with job scheduling systems such as SLURM on a supercomputer center. Therefore, another way to express multi-device parallelization is to employ the classical resource management system. SI Listing 6 demonstrates an example that submits a batch script to SLURM.

```
#!/bin/bash
#SBATCH -J <name>
#SBATCH -o <file name>
#SBATCH -p <str>
#SBATCH -n <int>
#SBATCH -N <int>
#SBATCH -c <int>

python your_script.py
```

**SI Listing 6** : The multi-device parallelization running of BrainPy models with the job scheduling system SLURM on a supercomputer. `your_scipt.py` defines the Python script to run your model.

#### 3.6 Customization of online and offline training methods

`brainpy.train.OnlineTrainer` and `brainpy.train.OfflineTrainer` supports to easily customize fitting methods by providing the `fit_method` parameter.

The customized offline training algorithm should follow the interface of `brainpy.algorithms.OfflineAlgorithm`, i.e., inheriting `brainpy.algorithms.OfflineAlgorithm` as a subclass and implementing `initialize()` function for training variable initialization and `call()` function for weight updating.

SI Listing 7 shows an example that using the `Lasso` model in `scikit-learn` package [26] to customize a LASSO regression training algorithm. The full source code of using this customized training interface to infer the Lorenz chaotic dynamics please check SI Listing 18.

```
import brainpy as bp
from sklearn.linear_model import Lasso

# using "Lasso" model in scikit-learn package
class LassoMethod(bp.algorithms.OfflineAlgorithm):
  def __init__(self, alpha=1., max_iter=int(1e4)):
    super(Lasso, self).__init__()
    self.model = Lasso(alpha=alpha, max_iter=max_iter)

  def call(self, identifier, target, input, predict=None):
    x = np.asarray(input)[0]
    y = np.asarray(target)[0]
    x_new = self.model.fit(x, y).coef_.T
    return bm.asarray(np.expand_dims(x_new, 1))

# initialize an offline trainer
trainer = bp.train.OfflineTrainer(
    <model>,
    fit_method=LassoMethod(),
    jit={'fit': False}
)
```

**SI Listing 7** : Using `Lasso` model in `scikit-learn` package to define an offline training algorithm. Since `Lasso` model is written in scikit-learn package, it is not compatible with the JIT compilation. Therefore, we turn off the JIT compilation in the `"fit"` phase.

SI Listing 8 presents a simple example that customizing an offline training algorithm by using BrainPy operators. The full code using this customized linear regression model to fit the chaotic dynamics is presented in SI Listing 19.

```
import brainpy as bp
import brainpy.math as bm

class LinearRegression(bp.train.OfflineAlgorithm):
  def __init__(self):
    super(LinearRegression, self).__init__()

  def call(self, identifier, targets, inputs, outputs=None):
    weights = bm.linalg.lstsq(inputs, targets)
    return weights[0]

# initialize an offline trainer
trainer = bp.train.OfflineTrainer(
    <model>,
    fit_method=LinearRegression()
)
```

**SI Listing 8** : Customizing a linear regression training algorithm.

Similarly, customization of a online training algorithm can be implemented by using the same way as the customization of a offline algorithm. Specifically, a new online training algorithm can be defined by subclassing `brainpy.train.OnlineAlgorithm`, in which `initialize()` function specifies how the training variables are initialized and `call()` function gives how the weight changes are computed accoording to the input, prediction and target data.

#### 3.7 Interoperation among BrainPy, JAX and NumPy

To make easy interoperability among BrainPy, JAX, and NumPy, BrainPy provides convenient interchanging interfaces. Transforming a NumPy's `ndarray` or JAX's `DeviceArray` as a `JaxArray` just needs to call `brainpy.math.asarray()` function. `JaxArray` can be transferred as a NumPy's `ndarray` through the build-in function `.to_numpy()` or `brainpy.math.as_ndarray()` function. Similarly, changing a `JaxArray` as a JAX's `DeviceArray` can utilize the `.to_jax()` build-in function or `brainpy.math.as_device_array()` function.

```
# changing a "ndarray" as a "JaxArray"
>>> brainpy.math.asarray(np.ones(2))
JaxArray([1., 1.], dtype=float32)

# changing a "JaxArray" as a "ndarray"
>>> brainpy.math.ones(2).to_numpy()
array([1., 1.], dtype=float32)

# changing a "DeviceArray" as a "JaxArray"
>>> brainpy.math.asarray(jnp.ones(2))
JaxArray([1., 1.], dtype=float32)

# changing a "JaxArray" as a "DeviceArray"
>>> brainpy.math.ones(2).to_jax()
DeviceArray([1., 1.], dtype=float32)
```

#### 3.8 Interoperation with other JAX frameworks

Many concepts of the object oriented programming interface of BrainPy are compatible with the functional programming of JAX (see main text). Therefore, BrainPy can be easily interoperated with other JAX frameworks.

In the below code snippet (SI Listing 9), we demonstrate that a `CNN` defined in Flax [97], a popular deep neural network library based on JAX, can be seamlessly integrated into a BrainPy program. Specifically, in the following code snippet, `CNN` is defined with Flax, then it is instantiated in `Network` class and used as a part of the whole model for feature extraction.

```
import brainpy as bp
import brainpy.math as bm
from jax.tree_util import tree_flatten, tree_map, tree_unflatten

from flax import linen as nn
import tensorflow_datasets as tfds

class CNN(nn.Module):
  """A CNN model implemented by Flax."""
  @nn.compact
  def __call__(self, x):
    x = nn.Conv(features=32, kernel_size=(3, 3))(x)
    x = nn.relu(x)
    x = nn.avg_pool(x, window_shape=(2, 2), strides=(2, 2))
    x = nn.Conv(features=64, kernel_size=(3, 3))(x)
    x = nn.relu(x)
    x = nn.avg_pool(x, window_shape=(2, 2), strides=(2, 2))
    x = x.reshape((x.shape[0], -1))
    x = nn.Dense(features=256)(x)
    x = nn.relu(x)
    return x

class Network(bp.dyn.DynamicalSystem):
  """A network model implemented by BrainPy"""
  def __init__(self):
    super(Network, self).__init__()

    # cnn and its parameters
    self.cnn = CNN()
    rng = bm.random.DEFAULT.split_key()
    params = self.cnn.init(rng, bm.ones([1, 4, 28, 1]).value)['params']
    leaves, self.tree = tree_flatten(params)
    self.implicit_vars.update(tree_map(bm.TrainVar, leaves))

    # rnn
    self.rnn = bp.layers.GRU(256, 100)

    # readout
    self.linear = bp.layers.Dense(100, 10)

  def update(self, sha, x):
    params = tree_unflatten(self.tree, [v.value for v in self.implicit_vars
     .values()])
    x = self.cnn.apply({'params': params}, bm.as_jax(x))
    x = self.rnn(sha, x)
    x = self.linear(sha, x)
    return x

def get_datasets():
  ds_builder = tfds.builder('mnist')
  ds_builder.download_and_prepare()
```

```
  train_ds = tfds.as_numpy(ds_builder.as_dataset(split='train', batch_size
    =-1))
  test_ds = tfds.as_numpy(ds_builder.as_dataset(split='test', batch_size
    =-1))
  train_ds['image'] = bm.float32(train_ds['image']) / 255.
  test_ds['image'] = bm.float32(test_ds['image']) / 255.
  return train_ds, test_ds

def loss_func(predictions, targets):
  logits = bm.max(predictions, axis=1)
  loss = bp.losses.cross_entropy_loss(logits, targets)
  accuracy = bm.mean(bm.argmax(logits, -1) == targets)
  return loss, {'accuracy': accuracy}

# get dataset
train_ds, _ = get_datasets()
X = bm.asarray(train_ds['image'].reshape((-1, 7, 4, 28, 1)))
Y = bm.asarray(train_ds['label'])

# initialize a network
net = Network()

# initialize a BPTT trainer
trainer = bp.train.BPTT(net,
                        loss_fun=loss_func,
                        optimizer=bp.optim.Momentum(0.1),
                        loss_has_aux=True)

# fitting the MNIST data
trainer.fit([X, Y], batch_size=100)
```

**SI Listing 9** : Integrating a CNN model defined with Flax into the BrainPy program.

### 4 Supporting Tables

**SI Table 1** : Caparison table of mathematical operators among NumPy [24], JAX [29], and BrainPy. The detailed comparison of the these operators please see https://brainpy.readthedocs.io/en/latest/apis/math/comparison.html. The number in each cell denotes the number of NumPy-like operators provided in each framework. The number in the bracket denotes the number of additional operators that are not provided in NumPy. BrainPy provides existing Numpy-like functions as much as possible, and also supports other special operator needs for brain dynamics programming.

| | Number of numerical functions | | |
|---|---|---|---|
| | in NumPy | in JAX | in BrainPy |
| Multi-dimensional Array | 56 | 42 | 46 (+8) |
| Array Operation | 399 | 313 | 341 (+11) |
| Linear Algebra Function | 19 | 19 | 19 |
| Discrete Fourier Transform | 18 | 18 | 18 |
| Random Number Generation | 51 | 16 | 48 (+8) |
| Dedicated Operator | - | - | (+20) |

**SI Table 2** : Numerical solvers provided in BrainPy for ordinary differential equations.

| Solver type | Solver name | Keyword |
|---|---|---|
| Runge-Kutta method | Euler | euler |
| | Midpoint | midpoint |
| | Heun's second-order method | heun2 |
| | Ralston's second-order method | ralston2 |
| | Second-order Runge-Kutta method | rk2 |
| | Third-order Runge-Kutta method | rk3 |
| | Four-order Runge-Kutta method | rk4 |
| | Heun's third-order method | heun3 |
| | Ralston's third-order method | ralston3 |
| | Third-order strong stability preserving Runge-Kutta method | ssprk3 |
| | Ralston's fourth-order method | ralston4 |
| | Fourth-order Runge-Kutta method with 3/8-rule | rk4_38rule |
| Adaptive Runge-Kutta method | Runge–Kutta–Fehlberg 4(5) | rkf45 |
| | Runge–Kutta–Fehlberg 1(2) | rkf12 |
| | Dormand–Prince method | rkdp |
| | Cash–Karp method | ck |
| | Bogacki–Shampine method | bs |
| | Heun–Euler method | heun_euler |
| Exponential method | Exponential Euler method | exp_euler |

**SI Table 3** : Numerical solvers provided in BrainPy for stochastic differential equations.

| Integral type | Solver name | Keyword | Scalar Wiener | Vector Wiener |
|---|---|---|---|---|
| Itô integral | Strong SRK scheme: SRI1W1 | srk1w1_scalar | Y | N |
| | Strong SRK scheme: SRI2W1 | srk2w1_scalar | Y | N |
| | Strong SRK scheme: KlPl | KlPl_scalar | Y | N |
| | Euler method | euler | Y | Y |
| | Milstein method | milstein | Y | Y |
| | Derivative-free Milstein method | milstein2 | Y | Y |
| | Exponential Euler | exp_euler | Y | Y |
| Stratonovich integral | Euler method | euler | Y | Y |
| | Heun method | heun | Y | Y |
| | Milstein method | milstein | Y | Y |
| | Derivative-free Milstein method | milstein2 | Y | Y |

**SI Table 4** : Numerical solvers provided in BrainPy for fractional differential equations.

| Derivative type | Solver name | Keyword |
|---|---|---|
| Caputo derivative | L1 schema | l1 |
| | Euler method | euler |
| Grünwald-Letnikov derivative | Short Memory Principle | short-memory |

### 5 Supporting Figures

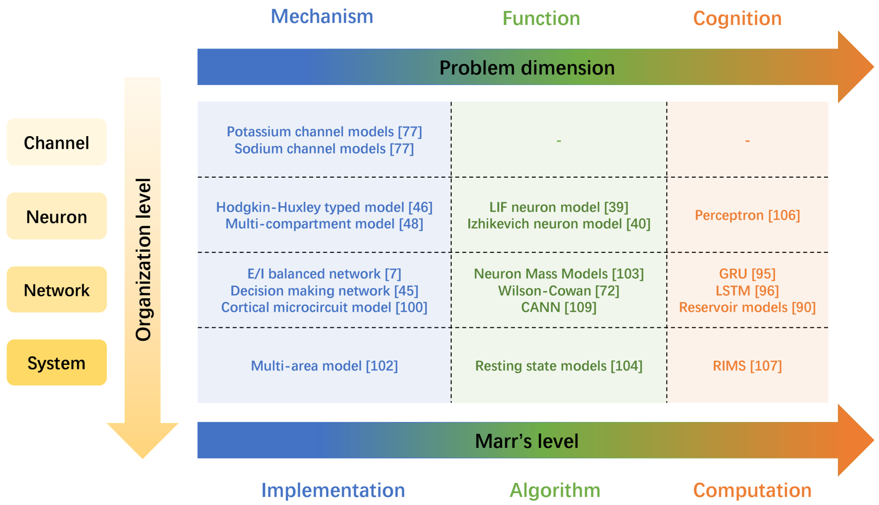

**SI Fig. 3** : **Summary of brain dynamics models across different organization levels and problem dimensions.** The *organization level* is the indicator for the anatomical scale of brain structure. Here, we choose organization levels commonly used in brain modeling, including *channel*, *neuron*, *network*, and *system*. The *problem dimension* and *Marr's level* are indicators for biophysical details. This summary is inspired by [98]. In each cell, we only list a few examples. The most biological grounded models are models at the **mechanistic implementation level**, which emphasize biological details for fine-grained simulations. Representatives of this kind of biophysical models are ion channel models [99], Hodgkin-Huxley typed neuron models [46], multi-compartment neuron models [48], spiking neural network models which mimic local populations [7, 45, 100], and large-scale system-level models which modeling functional systems [101, 102]. In contrast with realistic biophysical models, phenomenological models at the **functional algorithm level** focus on the qualitative behavior of the neural system. For instance, the reduced neuron models like LIF model [39] and Izhkevich model [40] take care of the spiking properties of neurons, rather than their ion channel dynamics. Similarly, population rate models like neural mass models [72, 75, 103] focus on the collective activity of neural population, rather than the single neuron dynamics. With these phenomenological models, ones build large-scale networks mimicking the whole brain [104, 105], in which each area is modeled as a simple population model, rather than large group of spiking neurons. Another category of brain dynamics modeling are data-driven models at the **cognitive computation level** which pay attention on high-level functions and reconstruct these functions through learning from data or tasks. In this sense, neurons are modeled as perceptron [106], and networks are modeled as recurrent neural networks [90, 95, 96, 107].

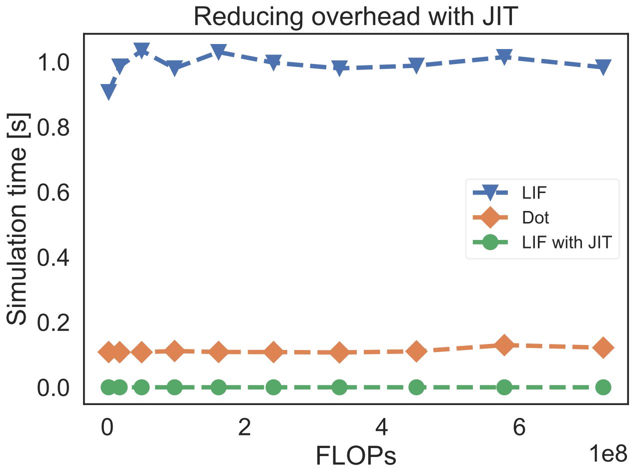

**SI Fig. 4** : **Reduce the overhead of a LIF model on the GPU device with JIT compilation**. Without JIT, the LIF model shows ten times slower than the matrix multiplication under the same FLOPs. After applying JIT compilation, the jitted LIF model shows much better performance than the matrix multiplication. This may result from the specific optimization of XLA on GPUs through operator fusion. The GPU device used here is NVIDIA Tesla V100.

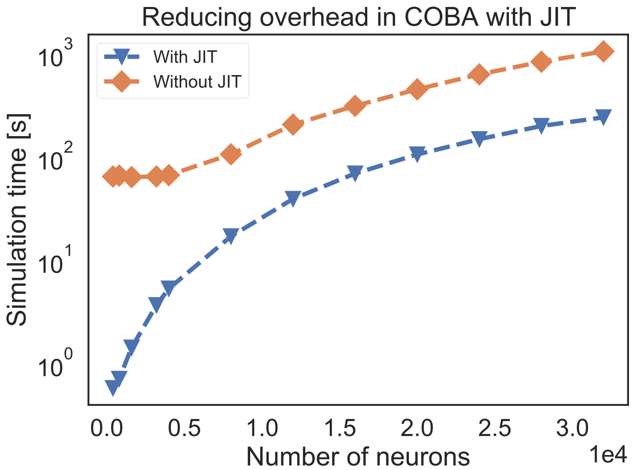

**SI Fig. 5** : **Reduce the overhead of a COBA network model [7] with JIT compilation**. After lowers the whole COBA network model into GPUs, the model obtains ten times of acceleration. The GPU device used here is NVIDIA Tesla V100.

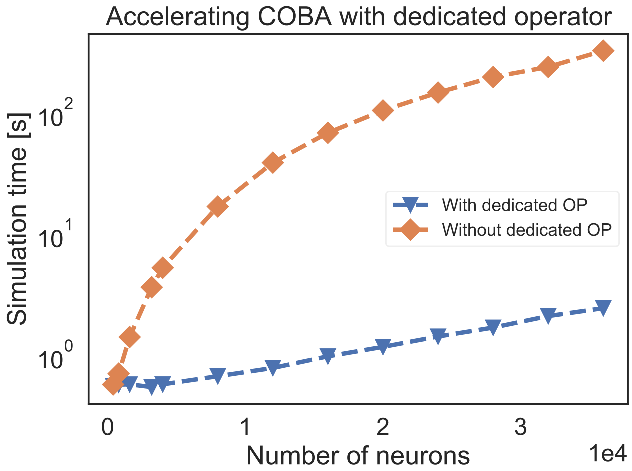

**SI Fig. 6 : Accelerating the COBA network model [7] with event-based primitive operators.** The acceleration effect is more prominent when the network size is large. The GPU device used here is NVIDIA Tesla V100.

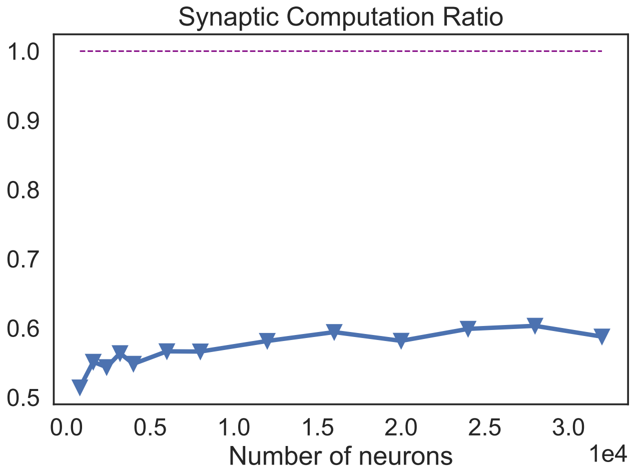

**SI Fig. 7 : The proportion of the time spent on the synaptic computation when simulating a COBA network model [7] with the event-based primitive operators.** This proportion is nearly half of the time of the whole network computation. It likely increases with the network size but in a mild way.

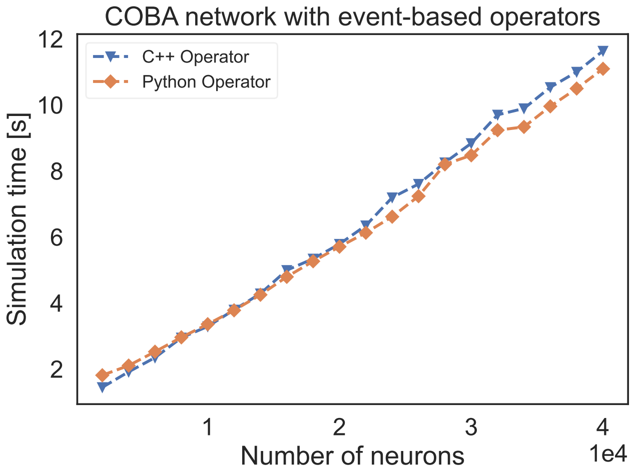

**SI Fig. 8** : **The speed comparison of event-based operators customized by C++ XLA custom call and our Python level registration interface**. "C++ Operator" presents the simulation time of a COBA network [44] using event-based operator coded by C++, and "Python Operator" shows the simulation speed of the network that implemented through our operator registered by the Python interface. The code for comparison please see SI Listing 20.

### 6 Supporting Codes

#### 6.1 Supporting codes for parallel simulations

```
import brainpy as bp
from joblib import Parallel, delayed, parallel_backend

class EINet(bp.dyn.Network):
  def __init__(self, scale=1.0):
    super(EINet, self).__init__()

    # network size
    num_exc = int(3200 * scale)
    num_inh = int(800 * scale)

    # neurons
    pars = dict(V_rest=-60., V_th=-50., V_reset=-60., tau=20., tau_ref=5.)
    self.E = bp.neurons.LIF(num_exc, **pars)
    self.I = bp.neurons.LIF(num_inh, **pars)

    # synapses
    prob = 0.02
    we = 0.6 / scale  # excitatory synaptic weight (voltage)
    wi = 6.7 / scale  # inhibitory synaptic weight
    self.E2E = bp.synapses.Exponential(self.E, self.E,
                  bp.conn.FixedProb(prob), g_max=we, tau=5.,
                  output=bp.synouts.COBA(E=0.))
    self.E2I = bp.synapses.Exponential(self.E, self.I,
                  bp.conn.FixedProb(prob), g_max=we, tau=5.,
                  output=bp.synouts.COBA(E=0.))
    self.I2E = bp.synapses.Exponential(self.I, self.E,
                  bp.conn.FixedProb(prob), g_max=wi, tau=10.,
                  output=bp.synouts.COBA(E=-80.))
    self.I2I = bp.synapses.Exponential(self.I, self.I,
                  bp.conn.FixedProb(prob), g_max=wi, tau=10.,
                  output=bp.synouts.COBA(E=-80.))

def run_ei_net(bg_current):
  net = EINet()
  runner = bp.dyn.DSRunner(
    net,
    monitors={'E.spike': net.E.spike},
    inputs=[(net.E.input, bg_current), (net.I.input, bg_current)],
    numpy_mon_after_run=False,
    progress_bar=False,
  )
  runner.run(100.)
  return runner.mon['E.spike']

if __name__ == '__main__':
  with parallel_backend(backend="threading"):
    parallel = Parallel(verbose=5)
    r = parallel([delayed(run_ei_net)(c) for c in [19., 20., 21., 22.]])
```

**SI Listing 10** : Multi-threading parallelization based on `joblib` for a E/I balanced network model, COBA [7].

```python
import brainpy as bp
import brainpy.math as bm
from jax import vmap

class EINet(bp.dyn.Network):
  def __init__(self, scale=1.0, method='exp_auto'):
    super(EINet, self).__init__()

    # network size
    num_exc = int(3200 * scale)
    num_inh = int(800 * scale)

    # neurons
    pars = dict(V_rest=-60., V_th=-50., V_reset=-60., tau=20., tau_ref=5.)
    self.E = bp.dyn.LIF(num_exc, **pars, method=method)
    self.I = bp.dyn.LIF(num_inh, **pars, method=method)

    # synapses
    prob = 0.02
    we = 0.6 / scale # excitatory synaptic weight (voltage)
    wi = 6.7 / scale  # inhibitory synaptic weight
    self.E2E = bp.synapses.Exponential(self.E, self.E,
                  bp.conn.FixedProb(prob), g_max=we, tau=5.,
                  output=bp.synouts.COBA(E=0.))
    self.E2I = bp.synapses.Exponential(self.E, self.I,
                  bp.conn.FixedProb(prob), g_max=we, tau=5.,
                  output=bp.synouts.COBA(E=0.))
    self.I2E = bp.synapses.Exponential(self.I, self.E,
                  bp.conn.FixedProb(prob), g_max=wi, tau=10.,
                  output=bp.synouts.COBA(E=-80.))
    self.I2I = bp.synapses.Exponential(self.I, self.I,
                  bp.conn.FixedProb(prob), g_max=wi, tau=10.,
                  output=bp.synouts.COBA(E=-80.))

def run_ei_net(bg_current):
  net = EINet()
  runner = bp.dyn.DSRunner(
    net,
    monitors={'E.spike': net.E.spike},
    inputs=[(net.E.input, bg_current), (net.I.input, bg_current)],
    numpy_mon_after_run=False,
    progress_bar=False,
  )
  runner.run(100.)
  return runner.mon['E.spike']

if __name__ == '__main__':
  r = vmap(run_ei_net)(bm.asarray([19., 20., 21., 22.]))
```

**SI Listing 11** : Multi-threading parallelization based on `jax.vmap` for a E/I balanced network model, COBA [7].

```
import brainpy as bp
from joblib import Parallel, delayed, parallel_backend

class EINet(bp.dyn.Network):
  def __init__(self, scale=1.0, method='exp_auto'):
    super(EINet, self).__init__()

    # network size
    num_exc = int(3200 * scale)
    num_inh = int(800 * scale)

    # neurons
    pars = dict(V_rest=-60., V_th=-50., V_reset=-60., tau=20., tau_ref=5.)
    self.E = bp.dyn.LIF(num_exc, **pars, method=method)
    self.I = bp.dyn.LIF(num_inh, **pars, method=method)

    # synapses
    prob = 0.02
    we = 0.6 / scale # excitatory synaptic weight (voltage)
    wi = 6.7 / scale  # inhibitory synaptic weight
    self.E2E = bp.synapses.Exponential(self.E, self.E,
                  bp.conn.FixedProb(prob), g_max=we, tau=5.,
                  output=bp.synouts.COBA(E=0.))
    self.E2I = bp.synapses.Exponential(self.E, self.I,
                  bp.conn.FixedProb(prob), g_max=we, tau=5.,
                  output=bp.synouts.COBA(E=0.))
    self.I2E = bp.synapses.Exponential(self.I, self.E,
                  bp.conn.FixedProb(prob), g_max=wi, tau=10.,
                  output=bp.synouts.COBA(E=-80.))
    self.I2I = bp.synapses.Exponential(self.I, self.I,
                  bp.conn.FixedProb(prob), g_max=wi, tau=10.,
                  output=bp.synouts.COBA(E=-80.))

def run_ei_net(bg_current):
  net = EINet()
  runner = bp.dyn.DSRunner(
    net,
    monitors={'E.spike': net.E.spike},
    inputs=[(net.E.input, bg_current), (net.I.input, bg_current)],
    numpy_mon_after_run=False,
    progress_bar=False,
  )
  runner.run(100.)
  return runner.mon['E.spike']

if __name__ == '__main__':
  with parallel_backend(backend="loky"):
    parallel = Parallel(verbose=5)
    r = parallel([delayed(run_ei_net)(c) for c in [19., 20., 21., 22.]])
```

**SI Listing 12** : Multi-processing parallelization based on `joblib` for a E/I balanced network model, COBA [7].

```python
import brainpy as bp
import brainpy.math as bm
from jax import pmap

class EINet(bp.dyn.Network):
  def __init__(self, scale=1.0, method='exp_auto'):
    super(EINet, self).__init__()

    # network size
    num_exc = int(3200 * scale)
    num_inh = int(800 * scale)

    # neurons
    pars = dict(V_rest=-60., V_th=-50., V_reset=-60., tau=20., tau_ref=5.)
    self.E = bp.dyn.LIF(num_exc, **pars, method=method)
    self.I = bp.dyn.LIF(num_inh, **pars, method=method)

    # synapses
    prob = 0.02
    we = 0.6 / scale # excitatory synaptic weight (voltage)
    wi = 6.7 / scale  # inhibitory synaptic weight
    self.E2E = bp.synapses.Exponential(self.E, self.E,
                  bp.conn.FixedProb(prob), g_max=we, tau=5.,
                  output=bp.synouts.COBA(E=0.))
    self.E2I = bp.synapses.Exponential(self.E, self.I,
                  bp.conn.FixedProb(prob), g_max=we, tau=5.,
                  output=bp.synouts.COBA(E=0.))
    self.I2E = bp.synapses.Exponential(self.I, self.E,
                  bp.conn.FixedProb(prob), g_max=wi, tau=10.,
                  output=bp.synouts.COBA(E=-80.))
    self.I2I = bp.synapses.Exponential(self.I, self.I,
                  bp.conn.FixedProb(prob), g_max=wi, tau=10.,
                  output=bp.synouts.COBA(E=-80.))

def run_ei_net(bg_current):
  net = EINet()
  runner = bp.dyn.DSRunner(
    net,
    monitors={'E.spike': net.E.spike},
    inputs=[(net.E.input, bg_current), (net.I.input, bg_current)],
    numpy_mon_after_run=False,
    progress_bar=False,
  )
  runner.run(100.)
  return runner.mon['E.spike']

if __name__ == '__main__':
  r = pmap(run_ei_net)(bm.asarray([19., 20., 21., 22.]))
  print(r.shape)
```

**SI Listing 13** : Multi-device parallelization based on `jax.pmap` for a E/I balanced network model, COBA [7].

#### 6.2 Supporting codes for dynamics analysis

```python
import brainpy as bp

bp.math.enable_x64()

gamma = 0.641  # Saturation factor for gating variable
tau = 0.06  # Synaptic time constant [sec]
a = 270.
b = 108.
d = 0.154
JE = 0.3725  # self-coupling strength [nA]
JI = -0.1137  # cross-coupling strength [nA]
JAext = 0.00117  # Stimulus input strength [nA]
mu = 20.  # Stimulus firing rate [spikes/sec]
coh = 0.5  # Stimulus coherence [%]
Ib1 = 0.3297
Ib2 = 0.3297

@bp.odeint
def int_s1(s1, t, s2, coh=0.5, mu=20.):
  I1 = JE * s1 + JI * s2 + Ib1 + JAext * mu * (1. + coh)
  r1 = (a * I1 - b) / (1. - bm.exp(-d * (a * I1 - b)))
  return - s1 / tau + (1. - s1) * gamma * r1

@bp.odeint
def int_s2(s2, t, s1, coh=0.5, mu=20.):
  I2 = JE * s2 + JI * s1 + Ib2 + JAext * mu * (1. - coh)
  r2 = (a * I2 - b) / (1. - bm.exp(-d * (a * I2 - b)))
  return - s2 / tau + (1. - s2) * gamma * r2

analyzer = bp.analysis.PhasePlane2D(
    model=[int_s1, int_s2],
    target_vars={'s1': [0, 1], 's2': [0, 1]},
    resolutions=0.001,
)
analyzer.plot_vector_field()
analyzer.plot_nullcline(coords=dict(s2='s2-s1'),
                        x_style={'fmt': '-'},
                        y_style={'fmt': '-'})
analyzer.plot_fixed_point()
analyzer.show_figure()
```

**SI Listing 14** : Phase plane analysis of a rate-based decision-making model [108]

```python
import brainpy as bp

bp.math.enable_x64()

@bp.odeint
def int_V(V, t, w, Iext=0.):
  return V - V * V * V / 3 - w + Iext

@bp.odeint
def int_w(w, t, V, a=0.7, b=0.8, tau=12.5):
  return (V + a - b * w) / tau

analyzer = bp.analysis.Bifurcation2D(
    model=[int_V, int_w],
    target_vars={'V': [-3, 3], 'w': [-3., 3.]},
    target_pars={'Iext': [0., 2.]},
    resolutions={'Iext': 0.005},
)
analyzer.plot_bifurcation(num_rank=10)
analyzer.plot_limit_cycle_by_sim()
analyzer.show_figure()
```

**SI Listing 15** : Bifurcation analysis of co-dimension 1 for a FitzHugh-Nagumo model [56].

```python
import brainpy as bp

bp.math.enable_x64()

@bp.odeint
def int_V(V, t, w, Iext=0.):
  return V - V * V * V / 3 - w + Iext

@bp.odeint
def int_w(w, t, V, a=0.7, b=0.8, tau=12.5):
  return (V + a - b * w) / tau

analyzer = bp.analysis.Bifurcation2D(
    model=[int_V, int_w],
    target_vars=dict(V=[-3, 3], w=[-3., 3.]),
    target_pars=dict(a=[0.5, 1.], Iext=[0., 2.]),
    resolutions={'a': 0.005, 'Iext': 0.005},
)
analyzer.plot_bifurcation(num_rank=10)
analyzer.plot_limit_cycle_by_sim()
analyzer.show_figure()
```

**SI Listing 16** : Bifurcation analysis of co-dimension 2 for a FitzHugh-Nagumo model [56].

```
import brainpy as bp
import brainpy.math as bm
import numpy as np

class CANN1D(bp.dyn.NeuGroup):
  def __init__(self, num, tau=1., k=8.1, a=0.5, A=10., J0=4., z_min=-bm.pi,
      z_max=bm.pi):
    super(CANN1D, self).__init__(size=num)

    # parameters
    self.tau = tau  # The synaptic time constant
    self.k = k  # Degree of the rescaled inhibition
    self.a = a  # Half-width of the range of excitatory connections
    self.A = A  # Magnitude of the external input
    self.J0 = J0  # maximum connection value
    self.z_min = z_min
    self.z_max = z_max
    self.z_range = z_max - z_min
    self.x = bm.linspace(z_min, z_max, num)  # The encoded feature values
    self.rho = num / self.z_range  # The neural density
    self.dx = self.z_range / num  # The stimulus density
    self.conn_mat = self.make_conn(self.x)

    # variables
    self.u = bm.Variable(bm.zeros(num))
    self.input = bm.Variable(bm.zeros(num))

    # function
    self.integral = bp.odeint(self.derivative)

  def derivative(self, u, t, Iext):
    r1 = bm.square(u)
    r2 = 1.0 + self.k * bm.sum(r1)
    r = r1 / r2
    Irec = bm.dot(self.conn_mat, r)
    du = (-u + Irec + Iext) / self.tau
    return du

  def dist(self, d):
    d = bm.remainder(d, self.z_range)
    d = bm.where(d > 0.5 * self.z_range, d - self.z_range, d)
    return d

  def make_conn(self, x):
    x_left = bm.reshape(x, (-1, 1))
    x_right = bm.repeat(x.reshape((1, -1)), len(x), axis=0)
    d = self.dist(x_left - x_right)
    return self.J0 * bm.exp(-0.5 * bm.square(d / self.a)) / (bm.sqrt(2 * bm
     .pi) * self.a)

  def get_stimulus_by_pos(self, pos):
    return self.A * bm.exp(-0.25 * bm.square(self.dist(self.x - pos) / self
     .a))

  def update(self, tdi):
    self.u.value = self.integral(self.u, tdi.t, self.input, tdi.dt)
    self.input[:] = 0.

def visualize_fixed_points(fixed_points, ax, num=10, title=None):
  ids = np.arange(0, fixed_points.shape[0],
                  int(fixed_points.shape[0] / num))
  for i in ids:
    ax.plot(fixed_points[i])
  ax.set_xticks([])
  ax.set_yticks([])
  if title: ax.set_title(title)
```

```python
def find_fixed_points_with_gd(A=10., tol=1e-6):
  cann = CANN1D(num=512, k=0.1, A=A, a=0.5)

  # initialize candidate fixed points
  candidates = cann.get_stimulus_by_pos(bm.linspace(-bm.pi, bm.pi, 1000).
     reshape((-1, 1)))
  candidates += bm.random.normal(0., 0.1, candidates.shape)

  # finding true fixed points from the generated candidate points
  finder = bp.analysis.SlowPointFinder(f_cell=cann,
                    target_vars={'u': cann.u}, dt=1.)
  finder.find_fps_with_gd_method(
    candidates={'u': candidates},
    tolerance=tol,
    optimizer=bp.optim.Adam(lr=bp.optim.ExponentialDecay(0.1, 2, 0.999)),
    num_batch=200,
  )
  finder.filter_loss(1e-5)
  finder.keep_unique()

  # visualize the found fixed points
  fig, gs = bp.visualize.get_figure(1, 1, 4.5, 6.)
  ax = fig.add_subplot(gs[0, 0])
  visualize_fixed_points(finder.fixed_points['u'], ax)

  # linearization analysis of the first 12 points
  finder.compute_jacobians(finder._fixed_points['u'][:12], plot=True)
```

**SI Listing 17** : Fixed point analysis of a continuous-attractor neural network [109] and the linearization analysis around the found fixed points.

#### 6.3 Supporting codes for customization of training algorithms

```
import brainpy as bp
import brainpy.math as bm
import matplotlib.pyplot as plt
import numpy as np
from sklearn import linear_model

bm.enable_x64()

def get_subset(data, start, end):
  res = {'x': data['x'][start: end],
         'y': data['y'][start: end],
         'z': data['z'][start: end]}
  X = bm.hstack([res['x'], res['y']])
  X = X.reshape((1,) + X.shape)
  Y = res['z']
  Y = Y.reshape((1,) + Y.shape)
  return X, Y

def plot_lorenz(x, y, true_z, predict_z, linewidth=None):
  plt.rcParams.update({"font.size": 15})

  fig, gs = bp.visualize.get_figure(3, 1, 1.5, 6)
  t_all = t_warmup + t_train + t_test
  ts = np.arange(0, t_all, dt)

  ax1 = fig.add_subplot(gs[0, 0])
  ax1.plot(ts[num_warmup + num_train:num_warmup + num_train + num_test],
           x[num_warmup + num_train:num_warmup + num_train + num_test],
           color='b', linewidth=linewidth)
  ax1.set_ylabel('x')
  ax1.axes.xaxis.set_ticklabels([])
  ax1.axes.yaxis.set_ticklabels([])
  ax1.axes.set_ybound(-21., 21.)
  ax1.axes.set_xbound(t_warmup + t_train - .5, t_all + .5)
  ax1.set_title('Reservoir Model')
  ax1.spines['right'].set_color('none')
  ax1.spines['top'].set_color('none')
  ax1.set_xticks([])
  ax1.set_yticks([])

  # testing phase y
  ax2 = fig.add_subplot(gs[1, 0])
  ax2.plot(ts[num_warmup + num_train:num_warmup + num_train + num_test],
           y[num_warmup + num_train:num_warmup + num_train + num_test],
           color='b', linewidth=linewidth)
  ax2.set_ylabel('y')
  ax2.axes.xaxis.set_ticklabels([])
  ax2.axes.yaxis.set_ticklabels([])
  ax2.axes.set_ybound(-26., 26.)
  ax2.axes.set_xbound(t_warmup + t_train - .5, t_all + .5)
  ax2.spines['right'].set_color('none')
  ax2.spines['top'].set_color('none')
  ax2.set_xticks([])
  ax2.set_yticks([])

  # testing phose z
  ax3 = fig.add_subplot(gs[2, 0])
  ax3.plot(ts[num_warmup + num_train:num_warmup + num_train + num_test],
           true_z[num_warmup + num_train:num_warmup + num_train + num_test
    ],
           color='b', linewidth=linewidth)
```

```
  ax3.plot(ts[num_warmup + num_train:num_warmup + num_train + num_test],
           predict_z[num_warmup + num_train:num_warmup + num_train +
     num_test],
           color='r', linewidth=linewidth)
  ax3.set_ylabel('z')
  ax3.set_xlabel('Time [ms]')
  ax3.axes.yaxis.set_ticklabels([])
  ax3.axes.set_ybound(3., 48.)
  ax3.axes.set_xbound(t_warmup + t_train - .5, t_all + .5)
  ax3.set_xticks([])
  ax3.set_yticks([])
  ax3.spines['right'].set_color('none')
  ax3.spines['top'].set_color('none')

  # plt.savefig(f'Reservoir-training.png', dpi=1000, transparent=True)
  plt.show()

dt = 0.02
t_warmup = 10.  # ms
t_train = 2.  # ms
t_test = 50.  # ms
num_warmup = int(t_warmup / dt)  # warm up NVAR
num_train = int(t_train / dt)
num_test = int(t_test / dt)

# Datasets #
lorenz_series = bp.datasets.lorenz_series(t_warmup + t_train + t_test,
                                          dt=dt,
                                          inits={'x': 17.67715816276679,
                                                 'y': 12.931379185960404,
                                                 'z': 43.91404334248268})

X_warmup, Y_warmup = get_subset(lorenz_series, 0, num_warmup)
X_train, Y_train = get_subset(lorenz_series, num_warmup, num_warmup +
     num_train)
X_test, Y_test = get_subset(lorenz_series, 0, num_warmup + num_train +
     num_test)

# Model #
class NGRC(bp.dyn.DynamicalSystem):
  def __init__(self, num_in):
    super(NGRC, self).__init__()
    self.r = bp.layers.NVAR(num_in, delay=4, order=2, stride=5, mode=bp.
     modes.batching)
    self.o = bp.layers.Dense(self.r.num_out, 1, mode=bp.modes.training)

  def update(self, sha, x):
    return self.o(sha, self.r(sha, x))

# Offline Training Algorithm #
class Lasso(bp.algorithms.OfflineAlgorithm):
  def __init__(self, alpha=1., max_iter=int(1e4)):
    super(Lasso, self).__init__()
    self.model = linear_model.Lasso(alpha=alpha, max_iter=max_iter)

  def __call__(self, identifier, y, x, outs=None):
    x = np.asarray(x)[0]
    y = np.asarray(y)[0]
    x_new = self.model.fit(x, y).coef_.T
    return bm.asarray(np.expand_dims(x_new, 1))

# Training the model
model = NGRC(2)
trainer = bp.train.OfflineTrainer(
```

```
    model,
    fit_method=Lasso(),
    jit={bp.running.FIT_PHASE: False}
)

# warm-up
outputs = trainer.predict(X_warmup)
print('Warmup NMS: ', bp.losses.mean_squared_error(outputs, Y_warmup))

# training
trainer.fit([X_train, Y_train])

# prediction
outputs = trainer.predict(X_test, reset_state=True)
print('Prediction NMS: ', bp.losses.mean_squared_error(outputs, Y_test))

# visualize the results
plot_lorenz(x=lorenz_series['x'].flatten().numpy(),
          y=lorenz_series['y'].flatten().numpy(),
          true_z=lorenz_series['z'].flatten().numpy(),
          predict_z=outputs.numpy().flatten())
```

**SI Listing 18** : Using a customized Lasso algorithm to train a reservoir computing model.

```
import brainpy as bp
import brainpy.math as bm
import matplotlib.pyplot as plt
import numpy as np
from sklearn import linear_model

bm.enable_x64()

def get_subset(data, start, end):
  res = {'x': data['x'][start: end],
         'y': data['y'][start: end],
         'z': data['z'][start: end]}
  X = bm.hstack([res['x'], res['y']])
  X = X.reshape((1,) + X.shape)
  Y = res['z']
  Y = Y.reshape((1,) + Y.shape)
  return X, Y

def plot_lorenz(x, y, true_z, predict_z, linewidth=None):
  plt.rcParams.update({"font.size": 15})

  fig, gs = bp.visualize.get_figure(3, 1, 1.5, 6)
  t_all = t_warmup + t_train + t_test
  ts = np.arange(0, t_all, dt)

  ax1 = fig.add_subplot(gs[0, 0])
  ax1.plot(ts[num_warmup + num_train:num_warmup + num_train + num_test],
           x[num_warmup + num_train:num_warmup + num_train + num_test],
           color='b', linewidth=linewidth)
  ax1.set_ylabel('x')
  ax1.axes.xaxis.set_ticklabels([])
  ax1.axes.yaxis.set_ticklabels([])
  ax1.axes.set_ybound(-21., 21.)
  ax1.axes.set_xbound(t_warmup + t_train - .5, t_all + .5)
  ax1.set_title('Reservoir Model')
  ax1.spines['right'].set_color('none')
  ax1.spines['top'].set_color('none')
  ax1.set_xticks([])
  ax1.set_yticks([])

  # testing phase y
  ax2 = fig.add_subplot(gs[1, 0])
  ax2.plot(ts[num_warmup + num_train:num_warmup + num_train + num_test],
           y[num_warmup + num_train:num_warmup + num_train + num_test],
           color='b', linewidth=linewidth)
  ax2.set_ylabel('y')
  ax2.axes.xaxis.set_ticklabels([])
  ax2.axes.yaxis.set_ticklabels([])
  ax2.axes.set_ybound(-26., 26.)
  ax2.axes.set_xbound(t_warmup + t_train - .5, t_all + .5)
  ax2.spines['right'].set_color('none')
  ax2.spines['top'].set_color('none')
  ax2.set_xticks([])
  ax2.set_yticks([])

  # testing phose z
  ax3 = fig.add_subplot(gs[2, 0])
  ax3.plot(ts[num_warmup + num_train:num_warmup + num_train + num_test],
           true_z[num_warmup + num_train:num_warmup + num_train + num_test
    ],
           color='b', linewidth=linewidth)
  ax3.plot(ts[num_warmup + num_train:num_warmup + num_train + num_test],
           predict_z[num_warmup + num_train:num_warmup + num_train +
    num_test],
           color='r', linewidth=linewidth)
```

```
  ax3.set_ylabel('z')
  ax3.set_xlabel('Time [ms]')
  ax3.axes.yaxis.set_ticklabels([])
  ax3.axes.set_ybound(3., 48.)
  ax3.axes.set_xbound(t_warmup + t_train - .5, t_all + .5)
  ax3.set_xticks([])
  ax3.set_yticks([])
  ax3.spines['right'].set_color('none')
  ax3.spines['top'].set_color('none')

  # plt.savefig(f'Reservoir-training.png', dpi=1000, transparent=True)
  plt.show()

dt = 0.02
t_warmup = 10.  # ms
t_train = 2.  # ms
t_test = 50.  # ms
num_warmup = int(t_warmup / dt)  # warm up NVAR
num_train = int(t_train / dt)
num_test = int(t_test / dt)

# Datasets #
lorenz_series = bp.datasets.lorenz_series(t_warmup + t_train + t_test, dt=
    dt,
                                          inits={'x': 17.67715816276679,
                                                 'y': 12.931379185960404,
                                                 'z': 43.91404334248268})

X_warmup, Y_warmup = get_subset(lorenz_series, 0, num_warmup)
X_train, Y_train = get_subset(lorenz_series, num_warmup, num_warmup +
    num_train)
X_test, Y_test = get_subset(lorenz_series, 0, num_warmup + num_train +
    num_test)

# Model #
class NGRC(bp.dyn.DynamicalSystem):
  def __init__(self, num_in):
    super(NGRC, self).__init__()
    self.r = bp.layers.NVAR(num_in, delay=4, order=2, stride=5, mode=bp.
    modes.batching)
    self.o = bp.layers.Dense(self.r.num_out, 1, mode=bp.modes.training)

  def update(self, sha, x):
    return self.o(sha, self.r(sha, x))

# Offline Training Algorithm #
class LinearRegression(bp.train.OfflineAlgorithm):
  def __init__(self):
    super(LinearRegression, self).__init__()

  def call(self, identifier, targets, inputs, outputs=None):
    weights = bm.linalg.lstsq(inputs, targets)
    return weights[0]

# Training the model
model = NGRC(2)
trainer = bp.train.OfflineTrainer(
    model,
    fit_method=LinearRegression(),
)

# warm-up
outputs = trainer.predict(X_warmup)
```

```
print('Warmup NMS: ', bp.losses.mean_squared_error(outputs, Y_warmup))

# training
trainer.fit([X_train, Y_train])

# prediction
outputs = trainer.predict(X_test, reset_state=True)
print('Prediction NMS: ', bp.losses.mean_squared_error(outputs, Y_test))

# visualize the results
plot_lorenz(x=lorenz_series['x'].flatten().numpy(),
            y=lorenz_series['y'].flatten().numpy(),
            true_z=lorenz_series['z'].flatten().numpy(),
            predict_z=outputs.numpy().flatten())
```

**SI Listing 19** : Using a customized linear regression algorithm to train a reservoir computing model.

#### 6.4 Supporting codes for customization of primitive operators

```
import brainpy as bp
import brainpy.math as bm

class EventSum(bm.XLACustomOp):
  """Customized operator."""

  def __init__(self):

    def abs_eval(events, indices, indptr, post_val, values):
      return post_val

    def con_compute(outs, ins):
      post_val = outs
      events, indices, indptr, _, values = ins
      for i in range(events.size):
        if events[i]:
          for j in range(indptr[i], indptr[i + 1]):
            index = indices[j]
            old_value = post_val[index]
            post_val[index] = values + old_value

    super(EventSum, self).__init__(abs_eval, con_compute)

# instantiate a customized op
event_sum = EventSum()

class ExponentialV1(bp.dyn.TwoEndConn):
  """Exponential synapse model using customized operator written in C++."""

  def __init__(self, pre, post, conn, g_max=1., delay=0., tau=8.0, E=0.):
    super(ExponentialV1, self).__init__(pre=pre, post=post, conn=conn)
    self.E = E
    self.tau = tau
    self.delay = delay
    self.g_max = g_max
    self.pre2post = self.conn.require('pre2post')
    self.g = bm.Variable(bm.zeros(self.post.num))
    self.integral = bp.odeint(lambda g, t: -g/self.tau, method='exp_auto')

  def update(self, tdi):
    self.g.value = self.integral(self.g, tdi.t, tdi.dt)
    self.g += bm.pre2post_event_sum(self.pre.spike, self.pre2post, self.
     post.num, self.g_max)
    self.post.input += self.g * (self.E - self.post.V)

class ExponentialV2(bp.dyn.TwoEndConn):
  """Exponential synapse model using customized operator written in C++."""

  def __init__(self, pre, post, conn, g_max=1., delay=0., tau=8.0, E=0.):
    super(ExponentialV2, self).__init__(pre=pre, post=post, conn=conn)
    self.E = E
    self.tau = tau
    self.delay = delay
    self.g_max = g_max
    self.pre2post = self.conn.require('pre2post')
    self.g = bm.Variable(bm.zeros(self.post.num))
    self.integral = bp.odeint(lambda g, t: -g/self.tau, method='exp_auto')

  def update(self, tdi):
```

```
    self.g.value = self.integral(self.g, tdi.t, tdi.dt)
    self.g += event_sum(self.pre.spike,
                        self.pre2post[0],
                        self.pre2post[1],
                        bm.zeros(self.post.num),
                        self.g_max)
    self.post.input += self.g * (self.E - self.post.V)

class EINet(bp.dyn.Network):
  def __init__(self, syn_type='v1'):
    syn_cls = ExponentialV1 if syn_type == 'v1' else ExponentialV2

    # neurons
    pars = dict(V_rest=-60., V_th=-50., V_reset=-60., tau=20., tau_ref=5.,
                V_initializer=bp.init.Normal(-55., 2.))
    E = bp.neurons.LIF(32000, **pars, method='exp_auto')
    I = bp.neurons.LIF(8000, **pars, method='exp_auto')

    # synapses
    E2E = syn_cls(E, E, bp.conn.FixedProb(0.02), E=0, g_max=0.06, tau=5)
    E2I = syn_cls(E, I, bp.conn.FixedProb(0.02), E=0, g_max=0.06, tau=5)
    I2E = syn_cls(I, E, bp.conn.FixedProb(0.02), E=-80, g_max=0.67, tau=10)
    I2I = syn_cls(I, I, bp.conn.FixedProb(0.02), E=-80, g_max=0.67, tau=10)

    super(EINet, self).__init__(E2E, E2I, I2E, I2I, E=E, I=I)

# network using the C++ operator
net1 = EINet(syn_type='v1')
runner1 = bp.dyn.DSRunner(net1, inputs=[('E.input', 20.), ('I.input', 20.)
    ])
t, _ = runner1.predict(10000., eval_time=True)
print(t)

# network using the Python oeprator
net2 = EINet(syn_type='v2')
runner2 = bp.dyn.DSRunner(net2, inputs=[('E.input', 20.), ('I.input', 20.)
    ])
t, _ = runner2.predict(10000., eval_time=True)
print(t)
```

**SI Listing 20** : The code to compare the simulation speed of primitive operators customized by C++ and Python interface.
